# *bioRheoFlow* A Rheometric Platform for In Situ Biofilm Mechanics Under Controlled Flow Conditions

**DOI:** 10.64898/2026.09.29.755442

**Authors:** Vincenzo Ianniello, Steffen Geisel, Eleonora Secchi, Theo Tervoort, Jan Vermant

## Abstract

Biofilm mechanics depend strongly on growth history, hydration, nutrient availability, and interfacial attachment, yet conventional rheology requires harvesting and transfer that can perturb the as-grown structure. Here, we introduce *bioRheoFlow*, a 3D-printed rheometer-mounted platform for cultivation and mechanical characterization without sample transfer. The platform combines evaporation compensation with static or axial nutrient flow, and benchmark measurements validate its rheometric performance. Applied to *Pseudomonas putida* and *Pantoea sp*., *bioRheoFlow* reveals that biofilms cultivated *in situ* under axial flow exhibit substantially higher elastic moduli than conventionally grown and scraped samples. Despite similar linear-viscoelastic stress ranges, the species show distinct nonlinear responses: *P. putida* exhibits a lower crossover stress and more abrupt failure, whereas *Pantoea sp*. sustains larger stresses and deformations. Creep suggests contributions from bulk deformation and biofilm–substrate interactions. *bioRheoFlow* provides a controlled framework for probing growth-conditioned mechanics in living biofilms.

## 1 Introduction

Biofilms are structured, surface-associated microbial communities embedded in a self-produced matrix of extracellular polymeric substances (EPS) [1–4]. They can be viewed as living soft materials whose mechanical response arises from the coupled contributions of the EPS matrix, microbial cells, spatial architecture, hydration, and substrate attachment. The resulting properties allow biofilms to maintain cohesion, resist externally imposed stresses, and persist under changing environmental conditions [5, 6]. Rheology provides a framework for quantifying how biofilms deform, flow, and fail under mechanical loading [3, 7, 8]. Such measurements are relevant to processes including cohesive deformation, fragmentation, detachment, and the formation of filamentous streamers that can obstruct medical and industrial flow systems [9–11]. Mechanical characterization nevertheless remains challenging because biofilms are heterogeneous living systems whose structure and properties vary spatially, evolve over time, and differ between nominally identical biological replicates [8, 10].

For bulk rheological characterization, biofilms are commonly grown on agar at an air–solid interface, harvested by scraping, and transferred to a rotational rheometer equipped with a parallel-plate or cone–plate geometry. Harvesting, transfer, and gap setting can irreversibly alter the as-grown structure, substrate interactions, and measured mechanical response, thereby contributing to experimental variability [8, 12–23]. Siri et al. [24] showed that gentle homogenization before bulk shear rheology can reduce structural variability and improve reproducibility, but at the cost of disrupting the native biofilm architecture. Conventional measurements therefore involve a fundamental trade-off between reproducibility and preservation of the mechanically relevant, as-grown structure. Hydration presents an additional difficulty: evaporation from exposed free surfaces changes water content, solids concentration, and microstructure, thereby biasing the measured viscoelastic response [25, 26]. Solvent traps, oil layers, humidity control, and alternative geometries can reduce solvent loss [25, 27–30], but may perturb the sample or prevent continuous medium exchange. Long-duration measurements of living biofilms consequently require simultaneous control of hydration, nutrient availability, and the surrounding chemical environment.

Several approaches have been developed to reduce sample disturbance and improve environmental control during biofilm rheology. Pavlovsky et al. [31] cultivated and characterized biofilms directly within a fixed-gap rheometer geometry. Waters et al. [32] similarly grew *Streptococcus mutans* biofilms directly on a rheometer plate within a controlled bioreactor and characterized their mechanical response *in situ*. Yannarell et al. [33] reduced disruption by transferring the biofilm together with its supporting membrane, while Kretschmer et al. [15] retained the growth substrate to quantify adhesive and cohesive failure using normal-force measurements. Geisel et al. [8] cultivated biofilms under controlled flow in a millifluidic device and subsequently transferred the biofilm-bearing substrate to a rotational rheometer. These approaches preserve different aspects of the as-grown state, but differ in whether cultivation and measurement occur in the same geometry, whether transfer remains necessary, and whether the method probes bulk shear rheology or substrate-mediated failure. A broadly applicable platform should therefore combine controlled hydration and nutrient delivery with minimal sample handling and compatibility with standard linear and nonlinear rheological protocols.

Flow during cultivation is itself an important state variable because it couples hydrodynamic forcing to nutrient and metabolite transport. Depending on the flow geometry, magnitude, and biofilm properties, hydrodynamic stresses can reshape surface-attached colonies or promote filamentous streamer formation, particularly in regions of strong extensional deformation [11, 34–37]. Flow also affects biomass accumulation, spatial organization, signaling, and EPS production by regulating the transport of nutrients and metabolic products [36, 38–40]. Mechanical conditioning during growth can consequently modify the subsequent rheological response; in several systems, increased hydrodynamic shear has been associated with stiffer and more elastic biofilms [41, 42]. Growth-flow conditions must therefore be controlled and distinguished from the shear imposed during rheological testing in order to determine how cultivation history conditions the measured mechanics.

Here, we introduce *bioRheoFlow*, a modular add-on for commercial rotational rheometers that enables biofilms to be cultivated and mechanically characterized directly within the rheometer gap under controlled hydration and medium-flow conditions. The platform combines evaporation compensation with configurable static or axial medium flow, allowing continuous nutrient delivery and long-duration measurements without harvesting or transferring the biofilm. The distinguishing feature of *bioRheoFlow* is the integration, within a single rheometer-mounted platform, of controlled medium exchange, configurable static or axial flow, optical monitoring of biofilm development, and subsequent linear and nonlinear rheological characterization without removing the biofilm from its growth environment. Biofilms are grown on porous glass plates whose pore size is selected to limit bacterial penetration and promote growth within the rheometrically probed region. We validate the control of solvent content and the rheometric response of the platform using volatile aqueous samples and a benchmark viscoelastic material. We then grow and characterize *Pseudomonas putida* and *Pantoea* sp. biofilms directly in *bioRheoFlow* to assess how *in situ* cultivation under axial nutrient flow, compared with conventional growth, scraping, and transfer, affects linear viscoelasticity, nonlinear deformation, creep, failure, and recovery. These comparisons determine whether cultivation history produces distinct mechanical states and whether species-dependent differences extend from small-deformation stiffness to large-deformation failure and restructuring. The modular components are fabricated by fused deposition modeling and stereolithography, and the corresponding design files are provided in the Supporting Information to facilitate adaptation to different rheometer configurations. *bioRheoFlow* therefore provides a common framework for distinguishing growth-conditioned mechanical behavior from artifacts introduced by evaporation, nutrient depletion, and sample transfer.

## 2 Results

### 2.1 Validation of *bioRheoFlow*

Rheometric validation was performed using a PVA/borax solution formulated to exhibit viscoelastic moduli in the range anticipated for the biofilms. Figure 2 compares frequency sweeps obtained using *bioRheoFlow* with measurements performed using conventional 10 mm and 25 mm parallel-plate geometries and a roughened 20 mm parallel-plategeometry. Within the common operating range of the geometries, the measured moduli were in agreement within 5%, indicating that integration of the flow cell did not substantially alter the measured linear viscoelastic response. The moments of inertia of the different tools are reported in Table S1 of the Supporting Information.

In oscillatory rotational rheometry, reliable measurements are restricted to an operating window within which the material response exceeds the instrument’s minimum torque and angular-displacement limits while remaining below the onset of significant tool or sample inertia [43]. The limits shown in Fig. 2 were calculated for the 10 mm plate and for the rheometer shaft carrying the *bioRheoFlow* upper casing without the fluidic needles. The corresponding moments of inertia are provided in the Supporting Information.

Evaporation compensation was assessed using oscillatory time sweeps on the same PVA/borax solution. Measurements were performed using a conventional geometry with and without the manufacturer’s evaporation hood and using *bioRheoFlow* in what we call “static mode”, with make-up solvent supplied through the lower porous plate. Solvent loss produced a systematic drift in |*G*^*\**^| in the conventional geometries. Defining the stable measurement interval as the time before |*G*^*\**^| deviated by more than 10% from its initial value, this threshold was reached after 8.5 min with the instrument’s standard evaporation blocker and after 2.2 min without it. With the *bioRheoFlow* setup, continuous infusion at 150 µLh^*−*1^ maintained |*G*^*\**^| near 3165 Pa, with a 10% deviation occurring only after approximately 100 min (see Fig.3). Active solvent replenishment therefore extended the stable measurement window from less than 10 min with the conventional hood to approximately 100 min.

### 2.2 Linear rheology of scraped biofilms

After validating evaporation compensation using the PVA/borax reference material, we applied it to biofilm characterization, first to conventionally scraped biofilms. Figure 4 compares measurements obtained with a standard 25 mm plate–plate geometry and with *bioRheoFlow* in static mode (Fig. 1a) for both *P. putida* and *Pantoea sp*. biofilms. In the static configuration, medium was supplied through the lower porous plate only to compensate for solvent loss, without axial flow through the measuring gap.

**Figure 1.**
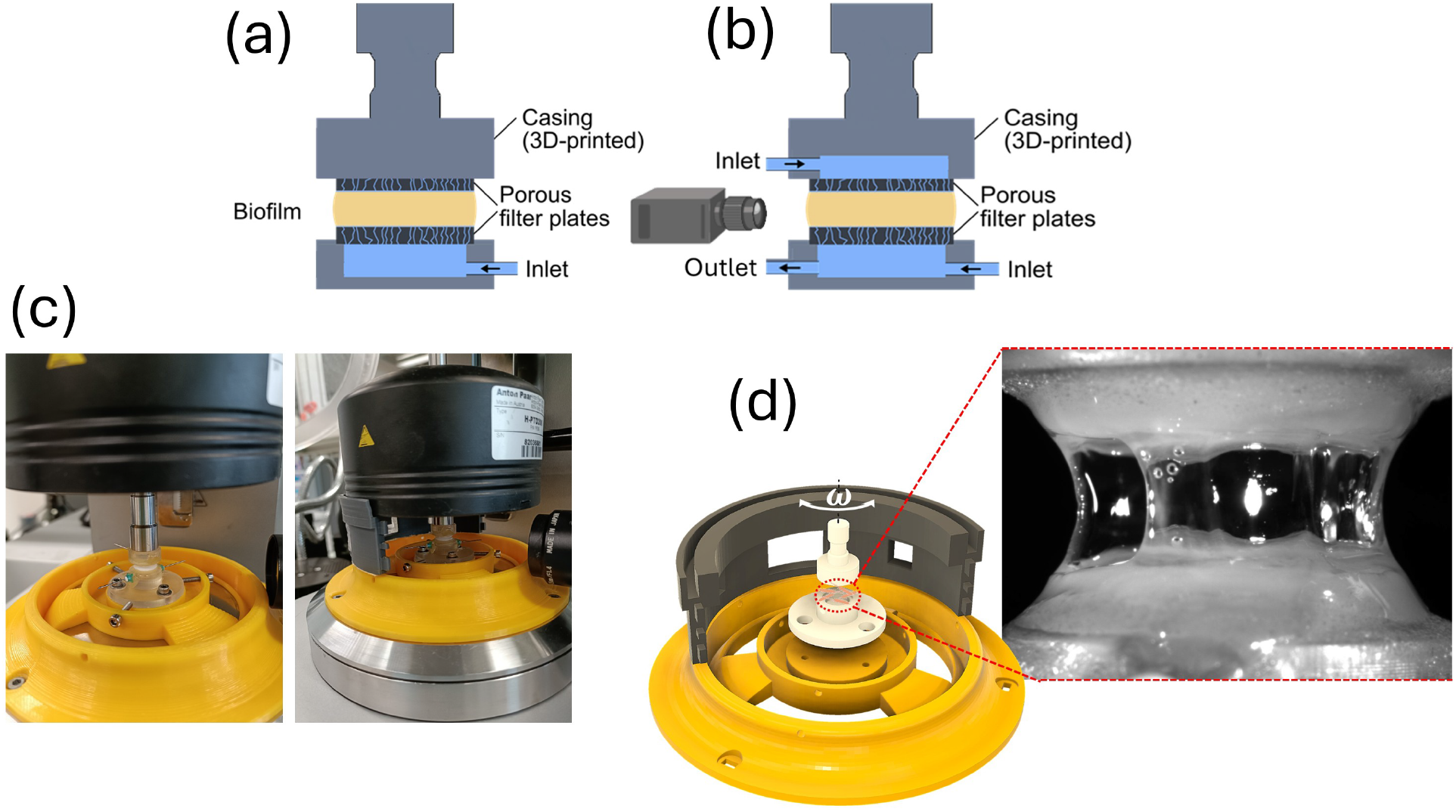
:Schematic and implementation of the *bioRheoFlow* platform. (a) Static configuration with a single bottom inlet. (b) Axial-flow configuration with two inlets (top and bottom) and one bottom outlet; the top inlet and outlet establish controlled axial flow within the measuring gap. (c) Photographs of the mounted platform on the rheometer. (d) CAD design of the device; inset shows a front view of the measuring gap where biofilm growth occurs.

**Figure 2.**
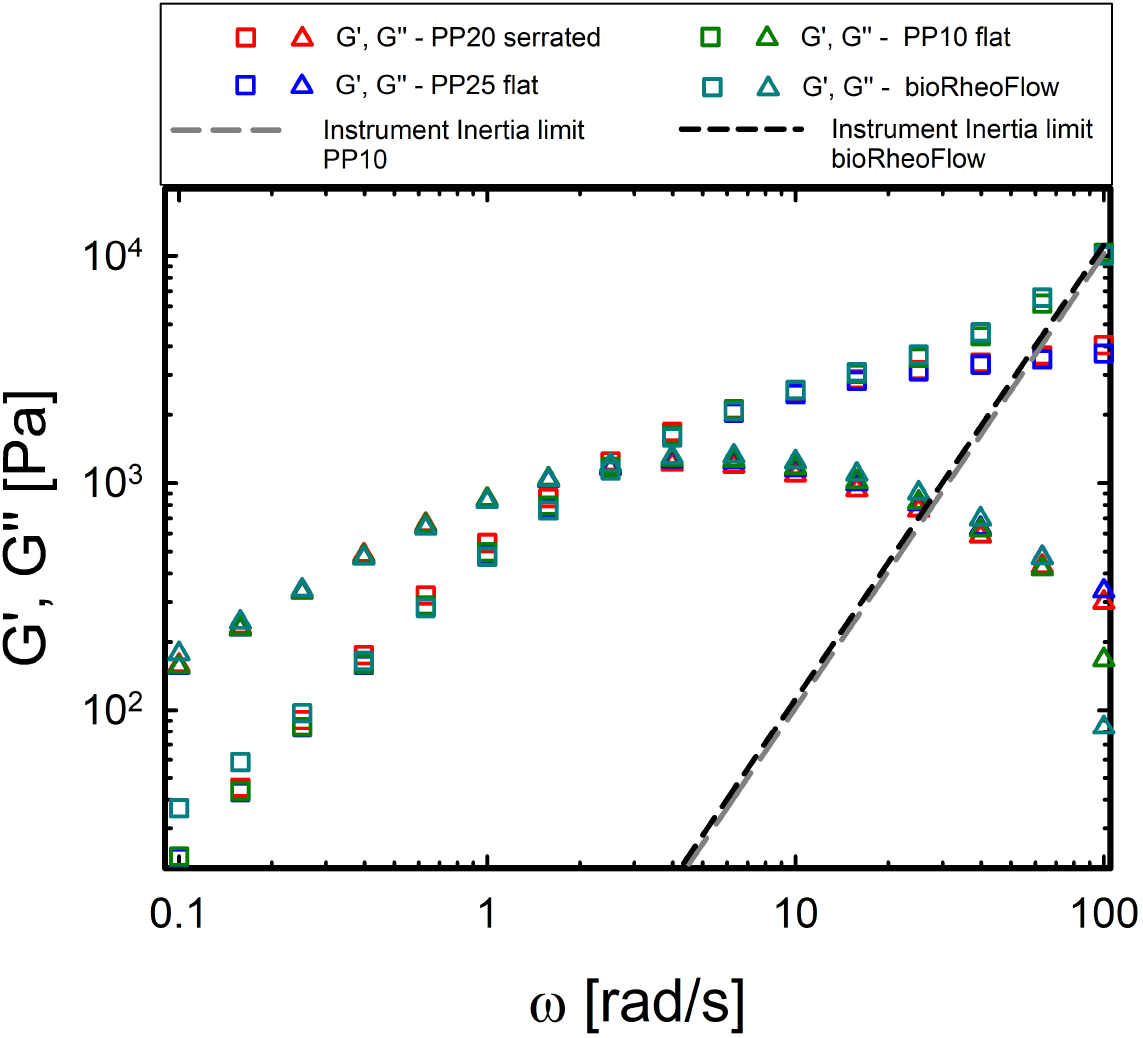
Oscillatory frequency sweeps obtained with different geometries at 25 °C and a shear strain amplitude of *γ* = 1%. The tested material was a PVA/borax solution prepared by mixing 50 mL of a 4 wt% aqueous PVA stock solution (*M*_*w*_ = 130 kDa) with 10 mL of a 4 wt% aqueous borax stock solution. The dashed lines indicate the calculated inertia limits for the conventional 10 mm plate and the *bioRheoFlow* upper assembly. Minimum-torque and angular-displacement limits lie outside the plotted range and are omitted for clarity.

**Figure 3.**
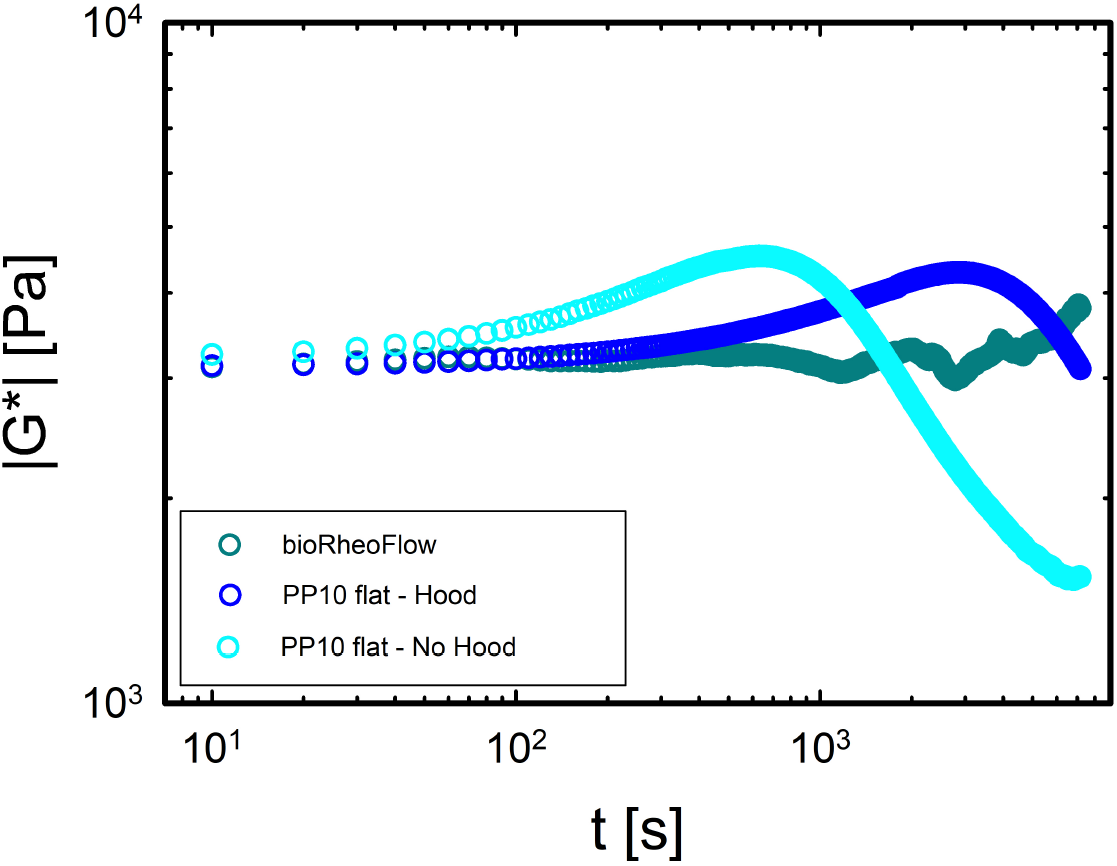
:Dynamic time sweep test conducted on PVA/Borax solution with a conventional plate–plate geometry and the proposed rheometric platform with continuous solvent flow. Measurements were conducted at a strain amplitude of *γ* = 1% and a frequency of *ω* = 10 rads^*−*1^.

**Figure 4.**
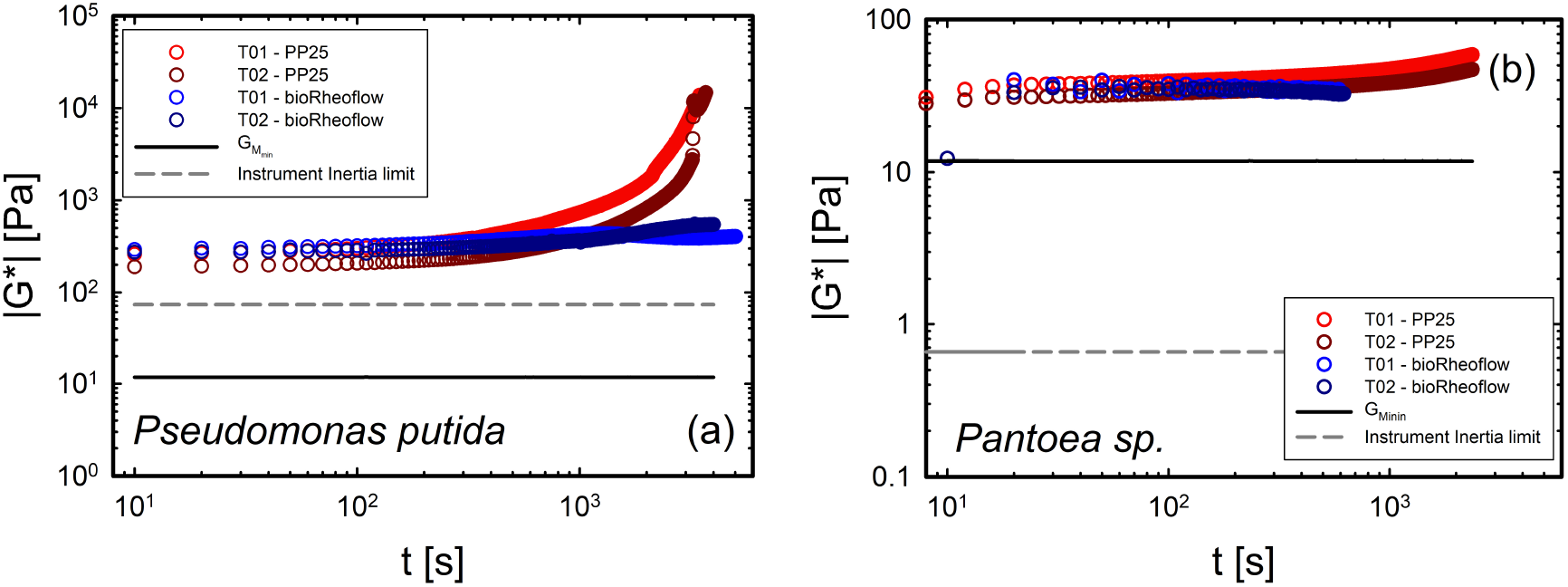
:Oscillatory time sweeps of scraped biofilms measured using a conventional 25 mm parallel-plate geometry and *bioRheoFlow* in static mode. (a) DTST of *P. putida* at *ω* = 10 rads^*−*1^. (b) DTST of *Pantoea sp*. at *ω* = 1 rads^*−*1^. The red symbols correspond to measurements performed with the conventional geometry, whereas the blue symbols correspond to measurements performed with the *bioRheoFlow* platform. Evaporation was compensated by solvent infusion at 50 µLh^*−*1^ for *P. putida* and 35 µLh^*−*1^ for *Pantoea sp*.

The lower frequency used for *Pantoea sp*. was selected because measurements at *ω* = 10 rads^*−*1^ were affected by the inertia limit of the measuring system, as indicated by the operational window in the corresponding frequency-sweep data (Fig. 5c). For the DTST comparison, conventionally scraped biofilms were measured using either a standard 25 mm parallel-plate geometry or *bioRheoFlow*, equipped with 10 mm-diameter porous glass measuring plates. For each species, the same frequency was used in both measuring configurations: *ω* = 10 rads^*−*1^ for *P. putida* and *ω* = 1 rads^*−*1^ for *Pantoea sp*.. Although the DTST measurements were therefore performed at different frequencies for the two species, the comparison remains meaningful because *G*^*′*^ of scraped *P. putida* is nearly frequency-independent over the relevant range, so its value at *ω* = 10 rads^*−*1^ is representative of that at *ω* = 1 rads^*−*1^.

**Figure 5.**
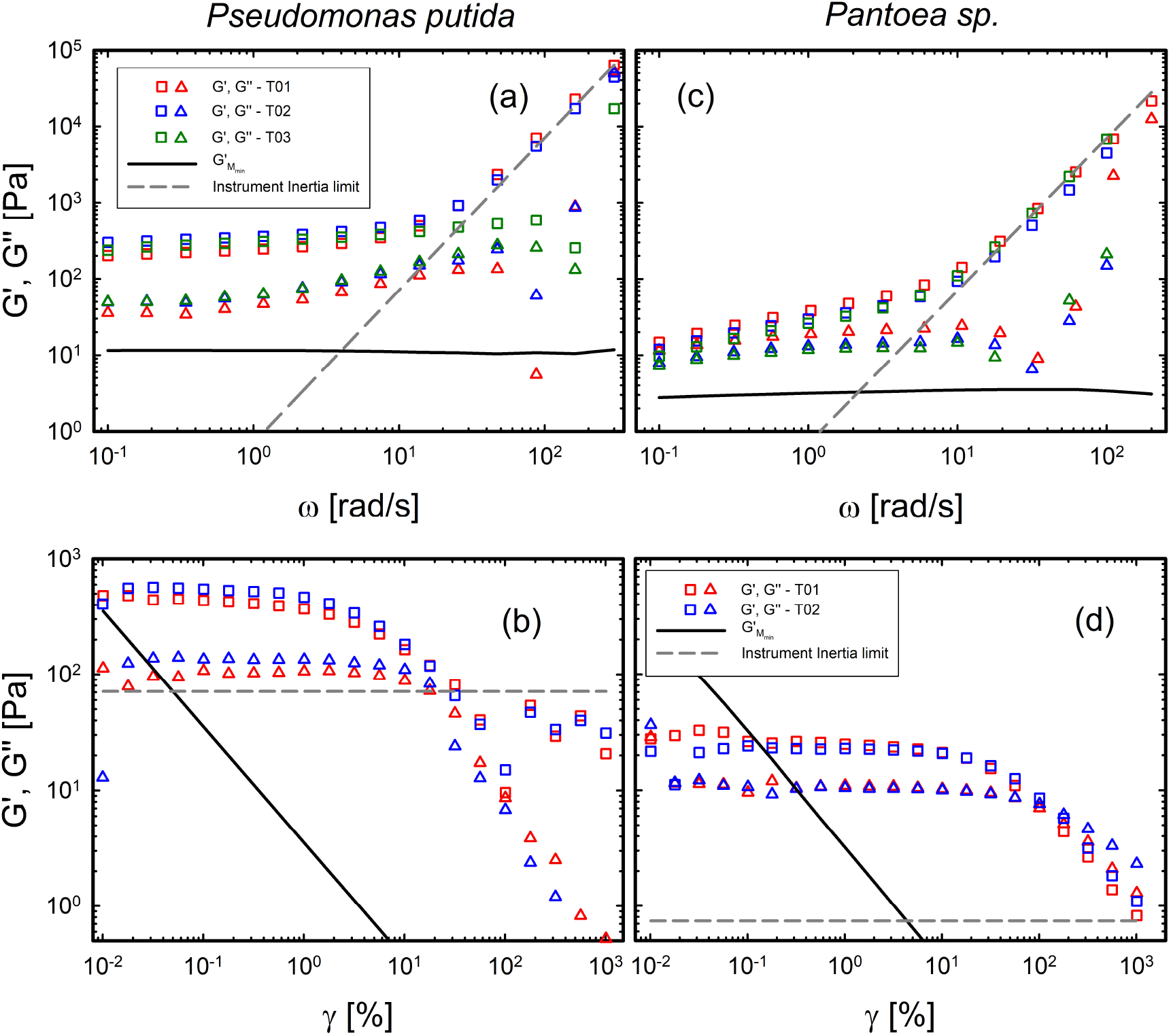
:DFST (a), (c) on scraped biofilms measured with the custom device for *P. putida* and *Pantoea sp*., respectively. DSST (b), (d) on scraped biofilms measured with the custom device for *P. putida* and *Pantoea sp*., respectively. The dashed line is the inertia limit according to the work of Ewoldt et al. [43].

In the conventional geometry, both biofilms exhibited systematic temporal drift despite the use of an evaporation hood (H-PTD 200, Anton Paar, Graz, Austria).

For scraped *P. putida*, the initial |*G*^*\**^| averaged approximately 270.7 Pa and deviated by 10% after approximately 1.67 min (Fig. 4a), while for *Pantoea sp*. the same deviation occurred after 4.33 min (average ∼ 37.5 Pa; Fig. 4b).

With continuous medium supply through *bioRheoFlow*, the temporal drift was reduced over the measurement interval. The corresponding mean values were approximately |*G*^*\**^| = 476 Pa for *P. putida* and |*G*^*\**^| = 37.4 Pa for *Pantoea sp*.. The close agreement between the two configurations for *Pantoea sp*. indicates that medium supply stabilized the measurement without substantially changing its initial modulus. In contrast, the higher modulus measured for *P. putida* with *bioRheoFlow* indicates sensitivity to geometry, mechanical deformation during loading, surface interactions, biological variability, or medium supply in addition to evaporation, pointing to the need for *in situ* growth to understand where this difference originates.

Figures 5a and 5c show the dynamic frequency sweep tests (DFST) for *P. putida* and *Pantoea sp*. biofilms, respectively. The *P. putida* biofilm exhibits a nearly frequency-independent storage modulus *G*^*′*^, consistent with a percolated elastic network characteristic of a strong gel. In contrast, *Pantoea sp*. shows a decrease in *G*^*′*^ toward lower frequencies, consistent with greater structural relaxation within the experimental window. These observations establish different rheological fingerprints but do not, by themselves, identify the molecular or microstructural origin of the responses.

The dynamic strain sweep tests (DSST) further demonstrate differences in stiffness and strain sensitivity (Figs. 5b and 5d). Within the linear viscoelastic regime, the plateau storage modulus was approximately *G*^*′*^ = 492.4 Pa for *P*.

*putida* and *G*^*′*^ = 24.4 Pa for *Pantoea sp*.. The critical strain, defined as the strain at which *G*^*′*^ decreases by 10% from its plateau value [44], is ∼ 1% for *P. putida* and ∼ 7.9% for *Pantoea sp*.. The procedure for determining the linear regime is detailed in the Supporting Information (Fig. S10). The crossover point (*G*^*′′*^ *> G*^*′*^), marking the onset of viscous-dominated behavior, occurs for *Pantoea sp*. at ∼ 139.8% strain with *G*^*′*^ = *G*^*′′*^ = 6.49 Pa. For *P. putida*, data above 20% strain are scattered, but extrapolation suggests a crossover near 30%.

Because the amplitude sweeps were performed at different frequencies, the critical and crossover strains should be interpreted as protocol-specific rather than directly comparable material constants. Overall, the measurements show that the scraped *P. putida* biofilm is stiffer and loses linearity at a lower strain, whereas the scraped *Pantoea sp*. biofilm is softer and remains linear over a larger strain range.

### 2.3 In situ growth under axial flow

*bioRheoFlow* enables biofilms to be cultivated directly within the measuring gap under axial nutrient flow (Fig. 1b). This protocol was compared with the conventional preparation in which biofilms were grown on agar, harvested by scraping, and measured using *bioRheoFlow* in evaporation-compensation mode (Fig. 1a). In the static evaporation-compensation configuration, medium was supplied through the lower porous plate to compensate solvent loss, whereas during *in situ* cultivation nutrient medium was continuously perfused through the measuring gap in the axial-flow configuration. Because these protocols differ in cultivation geometry, substrate and interfacial conditions, nutrient transport, and sample handling, the comparison assesses the combined influence of cultivation and preparation history rather than the effect of axial flow alone.

During *in situ* cultivation, the evolution of the rheometric signal was monitored for about 49 h for *P. putida* and 44 h for *Pantoea* sp. using an oscillatory time sweep at *ω* = 10 rads^*−*1^ and *γ* = 0.1% for both bacterial species (Fig. S8). A quasi-steady mechanical state was defined as a relative variation in |*G*^*\**^| below 10% over the final 2 h of cultivation, calculated from the change between the beginning and end of this interval relative to the value at its beginning. The corresponding variations were approximately 0.9% for *P. putida* and 1.7% for *Pantoea sp*., indicating a marked reduction in the rate of modulus evolution compared with the earlier stages of growth. Accordingly, the DTST was used as an operational indicator of biofilm development and stabilization rather than for quantitative determination of the biofilm moduli during axial flow.

After cultivation, axial nutrient flow was stopped and the platform was operated in static, evaporation-compensation mode for the subsequent frequency, strain-amplitude, and creep–recovery measurements. The upper geometry was lowered using the “viscoelastic” moving profile available in RheoCompass until complete filling of the measuring region was confirmed using the externally mounted camera aligned with the lateral optical window of *bioRheoFlow*. The resulting final measuring gap ranged from approximately 0.5 mm to 0.7 mm, depending on the initial inoculation gap and the extent of biofilm growth, according to the protocol described in Subsection 4.3.

The optical window permitted visual monitoring of the measuring gap during cultivation. Representative images of *P. putida* and *Pantoea sp*. biofilms grown under axial flow are provided in the Supporting Information.

The frequency sweeps (DFST) show that biofilms grown *in situ* under axial nutrient flow exhibit substantially higher elastic moduli than the conventionally grown and scraped counterparts (Figs. 6a,c). For *P. putida*, the plateau modulus increased from approximately 400 to 1000 Pa, whereas for *Pantoea sp. G*^*′*^ increased from approximately 24 to 380 Pa and developed a more pronounced low-frequency plateau. These results demonstrate that cultivation and preparation history strongly condition the measured linear viscoelastic response.

**Figure 6.**
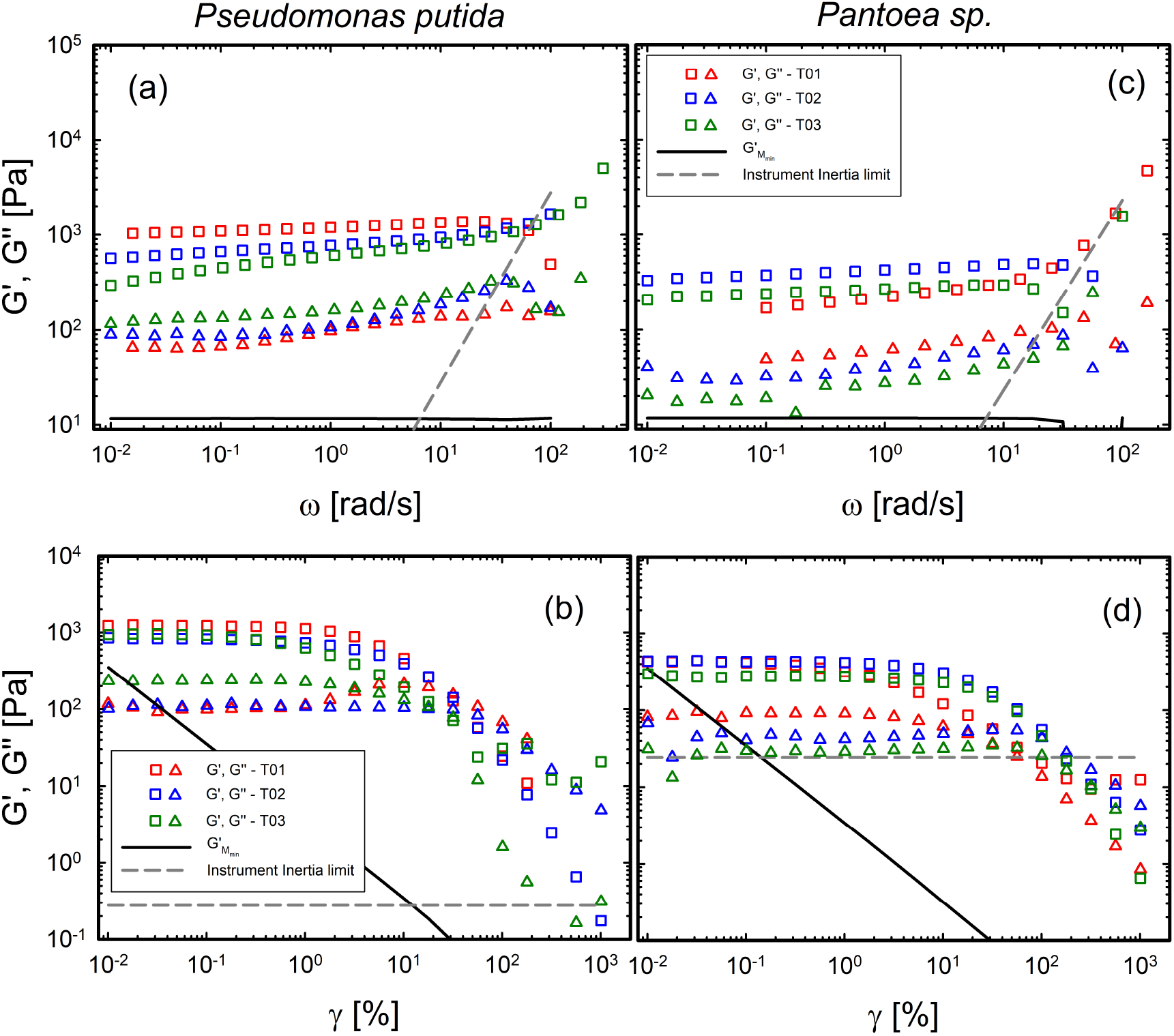
:DFST (a), (c) on biofilms grown *in situ* under axial nutrient flow for *P. putida* and *Pantoea sp*., respectively. DSST (b), (d) on biofilms grown *in situ* under axial nutrient flow for *P. putida* and *Pantoea sp*., respectively. According to the work of Ewoldt et al. [43], the solid and dashed lines refer to the minimum torque and inertia limits, respectively.

The strain-amplitude sweeps (DSST, Fig. 6b,d) show that the higher elastic moduli of biofilms grown under axial-flow conditions are accompanied by changes in their nonlinear response. Defining the critical strain, *γ*_*c*_, as the strain at which *G*^*′*^ decreases by 10% from its small-amplitude plateau, we obtain *γ*_*c*_ = 1.04 ±0.74% for *P. putida* and *γ*_*c*_ = 3.13 ±2.54% for *Pantoea sp*. (mean ± standard deviation, *n* = 3 independent biological replicates). The substantial variability, particularly for *Pantoea sp*., indicates that these values should be interpreted cautiously. The crossover (*G*^*′′*^ = *G*^*′*^), corresponding to a transition from predominantly elastic to predominantly dissipative behavior under the applied oscillatory protocol, occurs at a strain amplitude of approximately 35% for *P. putida* and 305% for *Pantoea sp*.. These crossover strains are protocol-dependent and should not be interpreted as unique yielding or flow-onset criteria.

Creep tests were used to further characterize the biofilms grown *in situ* under axial flow (Fig. 7). For *P. putida* (Fig. 7a), creep-compliance curves obtained at different stresses overlap up to *τ* = 5 Pa, indicating linear viscoelastic behavior; at 10 Pa the compliance increases relative to the lower-stress curves, indicating the onset of nonlinear deformation. For *Pantoea sp*. (Fig. 7b and Supplementary Fig. S12), the creep response starts to deviate from the stress-independent reference at approximately *τ* = 10 Pa, whereas the pronounced increase in compliance occurs only at substantially higher stresses, around *τ* = 80 Pa.

**Figure 7.**
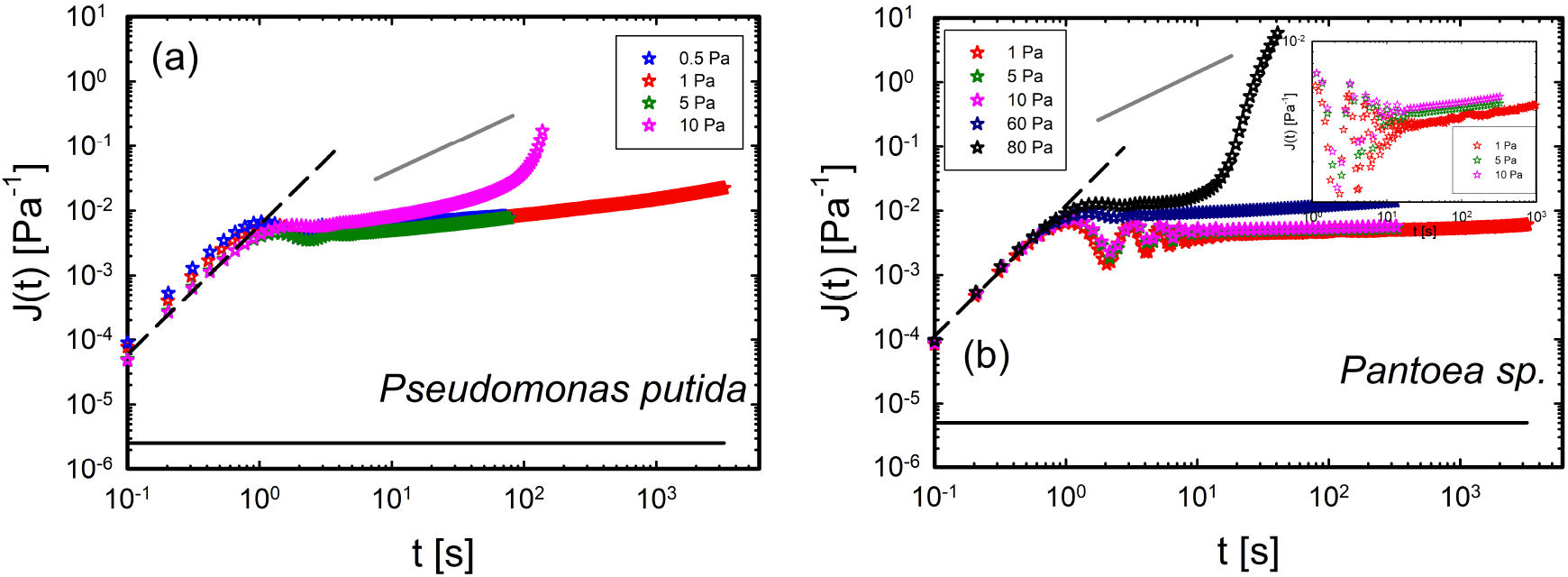
:Creep compliance of biofilms grown *in situ* under axial nutrient flow: (a) *P. putida*; (b) *Pantoea sp*.. Insets show the response at applied stresses of 1, 5, and 10 Pa. The dark gray line indicates unit slope (*∼ t*^1^), corresponding to the terminal flow regime. The dark solid lines correspond to the minimum compliance measurable by the rheometer (see Supporting Information). The dashed black lines indicate the acceleration of the motor (*∼ t*^2^).

For *Pantoea sp*., closer inspection reveals that the onset of nonlinear deformation occurs well below the stress associated with the subsequent abrupt increase in compliance, approximately 80 Pa (see inset in Fig. 7b). The creep curves at 1 Pa and 5 Pa nearly superimpose, consistent with linear viscoelastic behavior, whereas the response at 10 Pa deviates systematically from this reference. We therefore identify approximately 10 Pa as the onset of nonlinear deformation under this creep protocol. The much larger increase in compliance observed at higher stresses represents a distinct regime of accelerating deformation and possible progressive structural damage.

To assess whether the creep protocol produced persistent changes in the measured mechanical response, oscillatory time sweeps (DTST) and frequency sweeps (DFST) were performed before and after creep. For creep tests performed within the linear viscoelastic regime, a recovery period of approximately 3 min was applied before the post-creep frequency sweep. Recovery of the linear response was accepted when the post-creep *G*^*′*^ differed by less than 10% from the corresponding pre-creep value; the same criterion was applied to both bacterial species. Creep tests producing biofilm breakdown were not evaluated using this short-term criterion and their recovery kinetics are discussed separately in Subsection 2.5.

This comparison indicates recovery of the measured linear viscoelastic response, although it does not exclude transient structural disruption during creep.

At short times, the apparent creep compliance shown in Fig. 7 increases approximately as *J*(*t*) ∝ *t*^2^, followed by damped oscillations before the material-dominated response becomes accessible. The quadratic regime results from acceleration of the rheometer tool immediately after application of the stress step and is an instrumental inertia artifact that is independent of the applied stress when expressed as apparent compliance [43]. The subsequent oscillations, commonly termed creep ringing, arise from coupling between the rotational inertia of the instrument and the elastic response of the sample.

The latter is consistent with the DFST measurements. In particular, the gel-like behavior of the *Pantoea sp*. biofilm, characterized by a nearly frequency-independent storage modulus *G*^*′*^, may result in a higher propensity for ringing events before the compliance stabilizes.

Creep ringing can be used to extract short-time viscoelastic information beyond the frequency range directly accessible through conventional oscillatory measurements [45]. The resulting material functions are, however, obtained through an inertia-coupled constitutive model and therefore provide a model-dependent consistency check rather than a direct measurement of the frequency response. We analyzed the early-time creep response of *P. putida* and *Pantoea sp*. at stresses within their respective linear viscoelastic regimes using a Jeffreys-type constitutive model coupled to the rotational inertia of the upper measuring system. Related phenomenological models have previously been applied to biofilm rheology [31, 46–48]. The inertia of the upper measuring system was obtained from the “measuring system inertia” parameter provided by the rheometer software for the corresponding tool configuration and was incorporated into the equation of motion; the values are reported in Table S1 of the Supporting Information. Model parameters were obtained by fitting the early-time creep response over 0.2 s to 7.1 s for *P. putida* and 0.1 s to 25.2 s for *Pantoea sp*. and were compared with *G*^*′*^(*ω*) and *G*^*′′*^(*ω*) measured independently by oscillatory frequency sweeps over their common reliable frequency range. The complete model formulation, fitting procedure, and definition of the *χ*^2^ objective function are provided in the Supporting Information.

The model fits are shown in Fig. 8. The inertia-coupled Jeffreys model captures the early-time creep response and pronounced ringing observed for *Pantoea sp*.. For *P. putida*, the ringing is less pronounced and the agreement is poorer, is less pronounced and the agreement between the model and the experimental response is poorer, consistent with the larger value of the fitting objective function. The elastic modulus associated with the Kelvin–Voigt spring was extracted from the fits and compared with the storage modulus obtained independently from oscillatory frequency sweeps over the corresponding frequency range. This comparison provides a model-dependent consistency check between the transient and oscillatory measurements. The parameters obtained from the fitting are reported in Table 1.

**Table 1:** Parameters obtained by applying the Jeffreys model. The inertial term present in the model is reported in Table S1 in the Supporting Information.

| | $G_1$ [Pa] | $\eta_1$ [Pa s] | $\eta_2$ [Pa s] | $\omega$ [rad/s] | $\chi^2$ |
| --- | --- | --- | --- | --- | --- |
| *P. putida* | $3.17 \times 10^2$ | 0.64 | $2.33 \times 10^3$ | 2.45 | $6.2 \times 10^{-3}$ |
| *Pantoea sp.* | $2.72 \times 10^2$ | 0.33 | $2.88 \times 10^4$ | 3.1 | $2.3 \times 10^{-3}$ |

**Table 2:** Values of 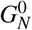 and *ξ* for the two bacterial species used in this work.

|  | *P. putida* |  | *Pantoea sp.* |  |
| --- | --- | --- | --- | --- |
|  | Scraped | Axial | Scraped | Axial |
| $G_N^0$ [Pa] | $492.34 \pm 84.93$ | $1004.87 \pm 206.42$ | $24.4 \pm 1.86$ | $381.2 \pm 88.49$ |
| $\xi$ [nm] | $25.3 \pm 1.5$ | $20.0 \pm 1.4$ | $68.9 \pm 1.8$ | $27.6 \pm 2.1$ |

**Figure 8.**
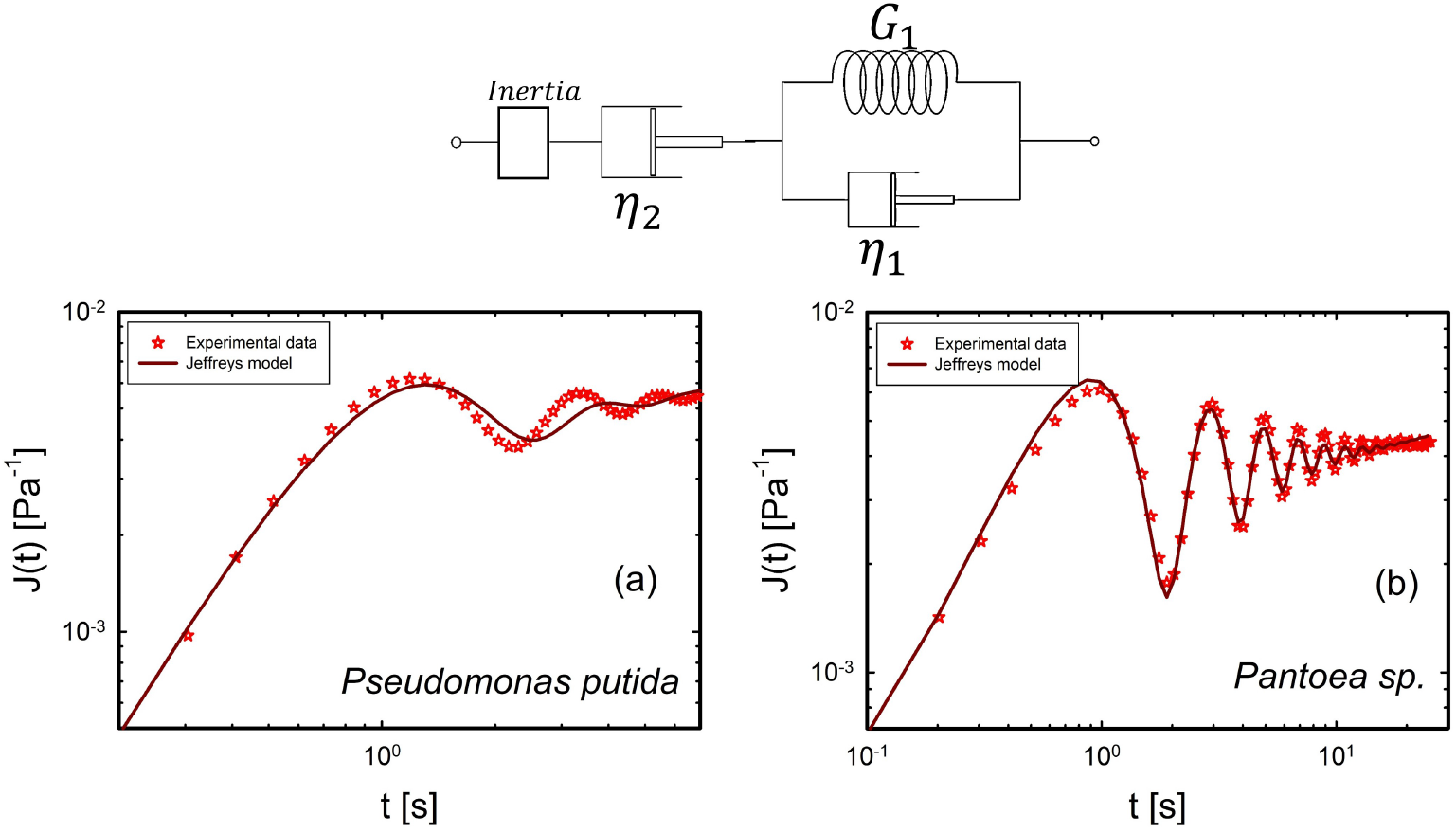
Scheme of the Jeffreys model. Fitting of the creep test at 1 Pa for (a) *P. putida* and (b) *Pantoea sp*.. The star symbols represent the experimental data, while the solid lines represent the model fitting.

As shown in Table 1, the elastic moduli obtained from the inertia-coupled fits were compared with the storage moduli measured independently by frequency sweeps over the corresponding frequency range. Because confidence intervals for the fitted parameters were not estimated, this comparison is interpreted as a model-dependent consistency check rather than as a quantitative test of agreement within experimental uncertainty. Furthermore, the parameter *η*_2_, which is associated with the steady-state flow viscosity, is strongly influenced by fitting over long time scales. For this reason, the analysis focuses only on the elastic transient response.

### 2.4 Evaluation of the network parameters

From the linear viscoelastic measurements, the elastic plateau modulus 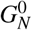 can be extracted. Following [49–51], 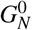 provides direct insight into the network microstructure, as it reflects the density of elastically active strands. In particular, the mesh size *ξ* , obtained here under a homogeneous-network approximation and defined as the average distance between crosslinks in the biofilm network, can be related to 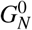 through Eq. 1.

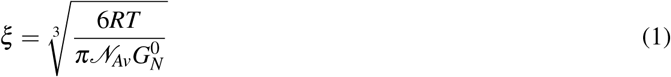

where *RT* represents the thermal energy, with 30 °C, and *N*_*Av*_ is the Avogadro’s number. The values of the elastic plateau modulus 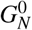 evaluated from bulk rheology and the average mesh size *ξ* from Eq. 1 are summarized in Tab. 2 for both biofilms according to the growth conditions.

### 2.5 Recovery after large deformation

We examined the recovery of biofilms grown *in situ* under axial flow after mechanical breakdown induced either by large-amplitude oscillatory deformation or by high-stress creep, to investigate if there could be long lived or irreversible changes. Oscillatory time sweeps show that *P. putida* re-establishes a stable storage modulus within approximately 8 hours whereas *Pantoea* sp. requires approximately 24 hours to recover its pre-deformation level (Figs. 9a,c). Following a second deformation cycle, the initially recovered modulus is lower, indicating incomplete or slower recovery at early times.

**Figure 9.**
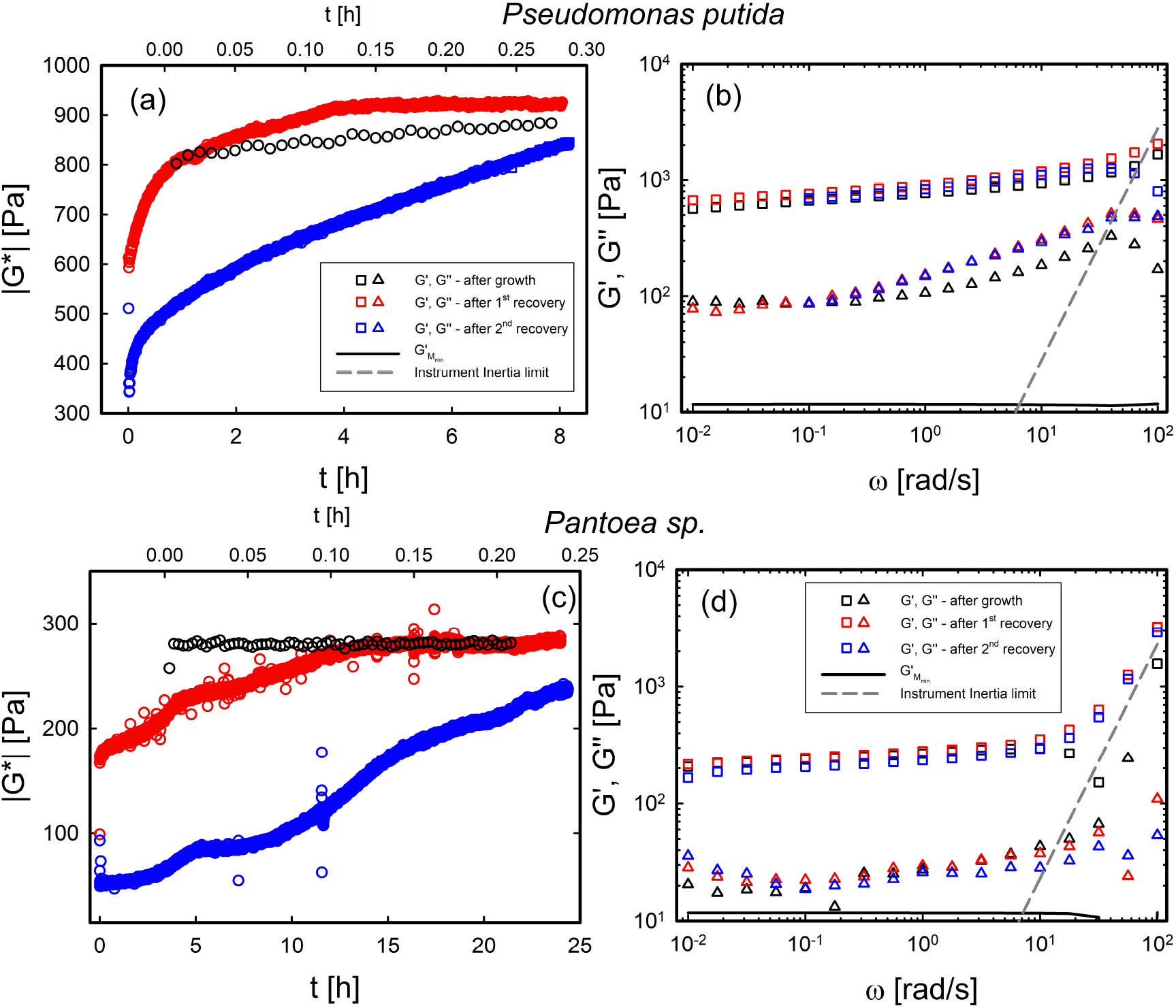
:(a, c) DTST on *P. putida* and *Pantoea sp*. biofilms after mechanical breakdown. The top x-axis refers to black symbols. (b, d) DFST on *P. putida* and *Pantoea sp*. after recovery. Black symbols indicate measurements after initial growth. In the DTST plots (a) and (c), the lines corresponding to the minimum detectable modulus and the instrument inertia limit lie outside the plotted axis range.

Comparison with the post-growth reference state (black symbols in Fig. 9) shows that both systems ultimately re-establish their viscoelastic properties, but time scales are very long. After a second disruption cycle, however, the initial modulus is reduced (blue symbols), suggesting partial loss of structural integrity during the early stages of structural recovery.

## 3 Discussion

We introduced and validated *bioRheoFlow*, a modular add-on for rotational rheometers that enables biofilm cultivation and mechanical characterization under controlled hydration and nutrient delivery. By combining evaporation compensation with static or axial medium flow, the platform supports long-duration measurements while minimizing changes caused by drying, nutrient depletion, harvesting, and transfer.

Biofilms cultivated *in situ* under axial nutrient flow were substantially stiffer than conventionally grown and scraped samples. For *P. putida*, the plateau storage modulus increased from approximately 400 to 1000 Pa. For *Pantoea sp*., *G*^*′*^ increased from 24.4 to ∼ 380 Pa, and developed a more pronounced elastic plateau. These differences reflect the combined effects of cultivation conditions, substrate attachment, and sample handling rather than axial flow alone.

Both biofilms departed from the linear viscoelastic regime at similar stresses of approximately 5–10 Pa, their nonlinear responses differed. *P. putida* was stiffer and showed a lower crossover stress and a more abrupt increase in creep compliance. *Pantoea* sp. sustained larger stresses and deformations before pronounced nonlinear deformation. The creep response likely includes contributions from both bulk deformation and the biofilm–substrate interface. Effective elastic length scales derived from the plateau moduli provide comparative measures of network connectivity, but should not be interpreted as direct geometric mesh sizes or evidence of uniform deformation.

At higher stresses, both systems exhibit regimes in which the logarithmic slope of the compliance exceeds unity, corresponding to accelerating creep rather than terminal viscous flow. This behavior is consistent with progressive structural damage or delayed failure. Because the measured response may include both deformation within the biofilm and changes at the biofilm–substrate interface, the present data do not distinguish uniquely among bulk fracture, interfacial slip, and partial detachment.

Comparison of DSST and creep data provides complementary information on the onset and progression of nonlinear deformation. In the strain-amplitude sweeps, both biofilms depart from the linear viscoelastic regime at stresses of approximately 5 Pa and 10 Pa for *P. putida* and *Pantoea sp*., respectively (see Fig. S11). The average crossover stress extracted from DSST is ∼30 Pa for *P. putida* and ∼60 Pa for *Pantoea sp*.. These values are of the same order as the stresses at which pronounced increases in creep compliance are observed (Fig. 7), although the two protocols probe nonlinear deformation differently and their characteristic stresses should not be expected to coincide exactly. Biofilms grown under axial flow therefore exhibit similar stresses for the onset of nonlinearity but distinct large-deformation responses. *P. putida* is stiffer and shows a more abrupt increase in compliance above the nonlinear threshold, consistent with a more strain-sensitive failure response. In contrast, *Pantoea sp*. sustains higher stresses and larger deformations before the pronounced compliance increase, indicating a more gradual and deformable nonlinear response.

For both biofilms, interpreting the plateau modulus using a homogeneous-network model yields an effective elastic length scale, *ξ* , smaller than the characteristic bacterial cell diameter, (*d*_*b*_ *≈* 1 µm). However, this estimate relies on an effective-medium interpretation of the plateau modulus, implicitly assuming a homogeneous polymer network. Such an assumption is likely oversimplified, because biofilms are structurally heterogeneous and exhibit deviations from linearity at very small strains (∼ 1%), indicative of pronounced strain localization rather than uniform affine deformation. Both systems depart from linear viscoelasticity at relatively small strains, indicating that their load-bearing structures are sensitive to deformation. Such early nonlinear softening is consistent with stress concentration at mechanically weak or heterogeneous regions, although the bulk rheological measurements do not establish strain localization directly. Differences in critical strain between cultivation conditions should therefore be interpreted as differences in deformation sensitivity rather than as evidence for a uniform affine network.

Within this framework, the confinement ratio *d*_*b*_*/ξ* should be interpreted cautiously. Rather than reflecting a uniform geometric constraint, it provides an effective measure of how cellular inclusions interact with a spatially heterogeneous matrix. The estimate is therefore useful as a comparative mechanical descriptor, but it cannot independently identify the underlying biofilm architecture or predict its failure stress.

From Table 3, *P. putida* exhibits a higher confinement number than *Pantoea sp*. under scraping conditions, whereas this difference becomes small under axial flow growth. However, under axial flow, *Pantoea sp*. sustains higher applied stresses before failure. This indicates that the confinement number alone does not directly predict the failure stress in these systems. The creep results therefore suggest that mechanical stability depends not only on the effective mesh size derived from 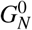, but also on structural heterogeneity and biofilm–substrate interactions. Any interpretation of confinement effects should thus be regarded as an effective description rather than evidence of a homogeneous network architecture.

**Table 3:** Calculated confinement number for *P. putida* and *Pantoea sp*.. Bacterial diameters used: *d*_*b*_ = 0.6 µm for *P. putida* and *d*_*b*_ = 0.8 µm for *Pantoea sp*.

|  | *P. putida* |  | *Pantoea sp.* |  |
| --- | --- | --- | --- | --- |
|  | Scraped | Axial | Scraped | Axial |
| $d_b/\xi$ | 23.7 | 30.0 | 11.6 | 29.0 |

These recovery measurements provide important context for comparing conventionally scraped biofilms with those grown *in situ* under axial nutrient flow. Scraping, transfer, and gap setting necessarily expose the biofilm to large and poorly controlled deformations that disrupt its as-grown architecture. Because recovery of the linear viscoelastic properties following a strong nonlinear deformation takes several hours, biofilms measured immediately after scraping are unlikely to have recovered from this deformation history. Their lower moduli may therefore reflect not only differences in cultivation conditions but also persistent mechanical disruption introduced during sample preparation.

In contrast, biofilms grown *in situ* remain attached to the measuring surfaces and are supplied continuously with medium, allowing hydration, nutrient availability, biological activity, and matrix remodeling to be maintained during growth. These conditions may also influence EPS composition. For *P. putida*, changes in cultivation conditions and water availability have been associated with shifts in the relative production of cellulose and alginate [52], while *Pantoea* sp. can form polysaccharide-rich extracellular matrices [53]. Such compositional differences could contribute to the observed changes in viscoelasticity, but matrix composition was not measured here. The higher moduli of the *in situ*-grown biofilms should therefore be interpreted as the combined consequence of growth conditions, preservation of the attached architecture, and avoidance of the large deformation associated with scraping and transfer.

Both biofilms recovered their linear viscoelastic response after large deformation, but very long recovery times were observed, on the order of hours. These experiments indicate that biofilms measured immediately after scraping and loading are unlikely to have recovered from the deformation imposed during sample preparation, confirming the relevance of the approach with in situ measurements developed here. Overall, *bioRheoFlow* provides a controlled framework for studying how cultivation history, structural heterogeneity, biological remodeling, and substrate interactions determine the time-dependent mechanics of living biofilms.

## 4 Methods

### 4.1 Bacterial culture and inoculation

The investigated biological systems are *P. putida* wild-type and *Pantoea sp*. wild-type. *P. putida* was grown overnight in 5 ml of lysogenic broth (LB), inoculated from frozen stock, at 30 °C while shaking at 30 rpm. Then, LB plates with 1% agar were inoculated from the overnight culture (100 µL). *Pantoea sp*. was grown overnight in Super Optimal Broth with Glucose (SOBG) medium, inoculated from frozen stock, at 30 °C while shaking at 30 rpm. Then, SOBG plates with 1.5% agar were inoculated from the overnight culture (100 µL). In both cases, the inoculum was evenly spread on the agar surface using a sterile L-shaped spreader. The plates were then incubated at 30 °C for 24 and 48 hours, respectively.

Agar-grown biofilms were then harvested by scraping and transferred into the rheometer using two glass microscopy slides. With a 10 mm measuring plate diameter, a single Petri dish, with a diameter of 9 cm, yields enough biofilm for up to five rheological replicates. No additional mechanical homogenization was applied. The scraped samples therefore experienced only the uncontrolled deformation associated with harvesting, transfer, and gap setting [24]. In contrast, for *in situ* cultivation within the flow cell, the bacterial suspension, taken from the same overnight culture, was directly inoculated into the rheometer using a syringe. Approximately 0.1 mL of bacterial suspension was introduced into the measuring gap. Detailed cultivation, gap-adjustment, flow, and rheological protocols are provided in Subsection 4.3.

### 4.2 *bioRheoFlow* design

The *bioRheoFlow* platform is a modular add-on designed to cultivate and mechanically characterize hydrated soft materials within the gap of a rotational rheometer (in this work an Anton Paar 302 was used). The assembly consists of upper and lower sample casings fabricated by stereolithography (Formlabs Form 3), a base (alignment) mount fabricated by fused deposition modeling (Prusa MK4), and an external cover that interfaces with the rheometer anti-evaporation hood. An overview of the assembled platform and its two flow configurations is shown in Fig. 1; detailed dimensions, fabrication parameters, and assembly instructions are provided in the Supporting Information (Fig. S1).

The casings accommodate porous glass filter plates (Ø 10 mm, height 2.8 mm; nominal pore size range 1 µm to 16 µm, Huberlab, Switzerland) bonded with Araldite (Huntsman, USA).

The plates provide rigid, rough growth surfaces intended to promote biofilm attachment and reduce interfacial slip while permitting medium transport. Their pore size was selected to limit bacterial penetration into the substrate. Because the porous plates form part of the mechanical and hydrodynamic boundary conditions, their permeability, roughness, and mounting procedure were kept constant between experiments.

The components were designed for additive manufacturing and adaptation through interchangeable mounts. The STL files, material specifications, printing parameters, and assembly procedure are provided in the Supporting Information.

The upper assembly mounts directly onto the Anton Paar disposable shaft, while the lower plate is fixed to a custom Peltier cover using a three-screw alignment system that ensures planarity and precise gap control (Fig. 1d). The procedure for assembling and aligning the tool prior to rheological measurements is described in the Supporting Information.

The external cover isolates the measuring system by enclosing the geometry within the manufacturer’s anti-evaporation hood, thereby limiting environmental exposure and contamination. It further enhances experimental control by integrating (i) a lateral optical window for real-time imaging using an externally mounted, vertically adjustable digital camera (Thorlabs Zelux CS165MU/M), featuring a 1440 *×* 1080 pixel CMOS sensor with a 3.45 *µ*m pixel pitch and an active area of 4.97 *×* 3.73 mm^2^ (1/2.9” format), operated in global shutter mode and coupled to a 25 mm focal length Navitar objective. (ii) opposing quartz halogen illuminator (Fiber-Lite 190, Dolan-Jenner) to improve visibility inside the chamber, and (iii) dedicated ports for tubing connections to the syringe pump (Standard PHD Ultra syringe pump, Harvard Apparatus), minimizing mechanical perturbations to the transducer.

The modular design is compatible with commercial rheometers and supports two flow configurations—static and axial—determined by inlet and outlet positioning (Fig. 1a,b). While developed for biofilms, the platform is equally suited for other hydrated or volatile soft materials, as demonstrated below.

#### Static Flow

The static configuration (Fig. 1a) provides continuous solvent or nutrient supply through the bottom plate to compensate evaporation and maintain hydration. Flow is introduced via a syringe tip glued beneath the porous glass filter and driven by a syringe pump (PHD Ultra, Harvard Apparatus) through 0.5 mm silicone tubing with an inner diameter of 0.5 mm, an outer diameter of 1.3 mm and a wall thickness of 0.4 mm (Huberlab, Switzerland). For biofilm measurements, this configuration preserves hydration during loading and testing while ensuring nutrient availability.

The infusion rates required to stabilize the viscoelastic response were 50 µLh^*−*1^ for *P. putida* and 35 µLh^*−*1^ for *Pantoea* sp., consistent with species-dependent water retention. After loading, the upper plate is lowered to the prescribed gap, excess material is trimmed using a spatula, and testing begins.

#### Axial Flow

In the axial configuration (Fig. 1b), a top inlet and a bottom outlet establish controlled axial nutrient medium flow through the measuring gap, with the configuration designed to distribute the axial medium flow uniformly across the measuring gap. A secondary bottom inlet enables transition to static conditions after growth without reversing flow, thereby avoiding reintroduction of spent medium and minimizing clogging risk. To prevent early bacterial washout, a thin (< 100 µm) agar layer was deposited on the lower porous plate. During growth, the lower auxiliary inlet remained closed. The inlet flow rate was set higher than the outlet flow rate to compensate for solvent loss by evaporation, following the same principle used in the static configuration. Optimal development occurred at inlet/outlet flow rates of 180*/*120 µLh^*−*1^ for *P. putida* and 140*/*90 µLh^*−*1^ for *Pantoea* sp. Growth was monitored optically until both filter surfaces were fully colonized (*≈* 48 h).

Because the upper flow connection introduces additional mechanical contributions to the measured signal due to the elastic response of the tubing and connections, quantitative rheology was performed only after switching back to the static configuration.

### 4.3 Biofilm rheological characterization protocol

The *bioRheoFlow* platform was mounted onto a conventional rotational rheometer (MCR 302, Anton Paar, Graz, Austria). To assess measurement reproducibility and enable direct comparison between bacterial species, growth conditions, all experiments were performed according to a standardized protocol.

All rheological measurements on biofilms were performed at 30 °C. Three independent biological replicates were investigated for each bacterial species and growth condition. For scraped biofilms, a single Petri dish provided sufficient material for up to five technical replicate measurements. Biofilms obtained from different Petri dishes were also compared to verify the reproducibility of the measured response. For measurements with the conventional configuration, scraped biofilms were characterized using a standard 25 mm parallel-plate geometry. For measurements with *bioRheoFlow*, both scraped and *in situ*-grown biofilms were characterized between the 10 mm-diameter porous glass plates integrated into the upper and lower sample casings. For biofilms grown *in situ*, each biological replicate corresponded to an independent inoculation and growth experiment. In the Results section, the measurements are presented as individual biological replicates.

Before starting the rheological measurements, the platform was disinfected with a 70:30 v/v solution of ethanol and distilled water for 30 minutes, followed by thorough flushing with growth medium to remove residual disinfectant. The platform was then flushed with 20 mL of the corresponding growth medium over approximately 20 min.

For conventionally scraped biofilms, the final measuring gap depended on the amount of loaded material and ranged from approximately 0.7 mm to 1.0 mm. After loading and gap setting, the sample was allowed to equilibrate until the normal force measured by the transducer decreased to 0 N, which generally required approximately 1 min.

Dynamic oscillatory time sweeps (DTST) were first performed on conventionally scraped biofilms to determine an appropriate medium-infusion rate for compensating solvent loss. Measurements were conducted at *ω* = 10 rads^*−*1^ for *P. putida* and *ω* = 1 rads^*−*1^ for *Pantoea sp*., using strain amplitudes of *γ* = 0.3% and *γ* = 1%, respectively, both within the linear viscoelastic regime, to monitor the time evolution of the moduli associated with solvent loss. The lower frequency used for *Pantoea sp*. was selected because measurements at *ω* = 10 rads^*−*1^ were affected by the inertia limit of the measuring system, as shown by the operational window reported in the DFST plots. Although different frequencies were used for the two species, comparison remained possible because the rheological response of scraped *P. putida* biofilms was nearly frequency-independent over the investigated frequency range.

Infusion rates ranging from 30 µLh^*−*1^ to 170 µLh^*−*1^ for *P. putida* and from 30 µLh^*−*1^ to 160 µLh^*−*1^ for *Pantoea sp*. were evaluated in preliminary screening tests. Rates that produced an evident non-stationary response were monitored for 10 min to 15 min, which was sufficient to identify either excessive or insufficient infusion. Excessively high infusion rates caused liquid accumulation within the measuring gap and visible dilution of the biofilm, whereas insufficient infusion resulted in a progressive change in the measured modulus and visible shrinkage of the biofilm between the plates. Biofilm volume changes were monitored using an externally mounted camera aligned with the lateral optical window of *bioRheoFlow*.

Because biofilm restructuring or recovery may also cause temporal evolution, modulus stability was interpreted as an operational criterion for evaporation compensation rather than as proof of an invariant biofilm structure. Evaporation compensation was operationally defined by a variation in the complex modulus |*G*^*\**^| of less than approximately 10% relative to its plateau value, together with the absence of visible biofilm shrinkage or liquid accumulation within the measuring gap. The lowest infusion rate ensuring stable moduli was therefore identified empirically. The selected infusion rates were 50 µLh^*−*1^ for *P. putida* and 35 µLh^*−*1^ for *Pantoea sp*.. At these selected infusion rates, each DTST was performed for a fixed duration of 100 min and was not interrupted before completion.

Subsequently, dynamic frequency sweeps (DFST) were conducted over *ω* = 100 rads^*−*1^ to *ω* = 0.1 rads^*−*1^. The sweeps were performed from high to low frequency using five measurement points per decade, with strain amplitudes of *γ* = 0.3% for *P. putida* and *γ* = 1% for *Pantoea sp*.. A waiting time of approximately 1 min was applied before each frequency sweep.

The linear viscoelastic range was verified through a dynamic strain sweep test (DSST) performed at the same frequency as the DTST, across a strain range of *γ* = 0.01–1000%. The strain amplitude was increased from low to high strain using four measurement points per decade. The upper boundary of the linear viscoelastic regime was defined as the strain at which *G*^*′*^ deviated by 10% from its initial plateau value.

For *in situ* growth measurements under axial flow, the bacterial suspension was inoculated at an initial measuring gap ranging from approximately 0.95 mm to 1.10 mm. The evolution of the rheometric response during biofilm growth was monitored through a DTST at *ω* = 10 rads^*−*1^ and *γ* = 0.1%. Because the axial-flow connection contributes to the measured mechanical response, the moduli recorded during cultivation were used to monitor their temporal evolution rather than as quantitative measures of the biofilm viscoelastic properties.

This small-amplitude oscillatory deformation was selected within the linear viscoelastic regime to monitor biofilm growth without significantly perturbing the developing network.

The cultivation period was approximately 48 h. A mature biofilm was operationally identified by stabilization of the rheometric signal together with optical confirmation that the biofilm filled the measuring gap and visibly contacted both porous plates. The optical assessment was performed using an externally mounted camera aligned with the lateral optical window of *bioRheoFlow*, and representative images are reported in the Supporting Information.

After growth, the upper geometry was lowered using the “viscoelastic” moving profile available in RheoCompass until complete filling of the measuring region was confirmed using an externally mounted camera aligned with the lateral optical window of *bioRheoFlow*. The resulting final measuring gap typically ranged from approximately 0.5 mm to 0.7 mm, depending on the initial inoculation gap and the extent of biofilm growth. The final position was selected based on optical confirmation of complete contact between the biofilm and both measuring surfaces; normal force was not used as a stopping criterion during this adjustment.

Before quantitative rheological measurements, the setup was switched from the axial-flow configuration to the static configuration. The linear and nonlinear mechanical response was then characterized through frequency sweeps, strain sweeps, and creep–recovery tests.

Creep tests were performed at increasing applied stresses to probe the transition from linear viscoelastic behavior to nonlinear deformation and breakdown. Oscillatory frequency sweeps were performed before and after creep to assess whether the applied stress produced persistent changes in the linear viscoelastic response. For creep tests performed within the linear viscoelastic regime, a recovery period of approximately 3 min was applied before the post-creep frequency sweep. Recovery was considered complete when the post-creep *G*^*′*^ differed by less than 10% from the corresponding pre-creep value. The same criterion was applied to both bacterial species. Tests producing biofilm breakdown were excluded from this short-term recovery criterion and were evaluated separately through long-duration recovery measurements, for which mechanical breakdown was induced either by a large-amplitude dynamic strain sweep or by creep at sufficiently high applied stress. Following breakdown, the evolution of the viscoelastic response was monitored through oscillatory measurements until a stable mechanical response was re-established.

Data analysis was performed using custom routines in MATLAB^®^ R2021a (MathWorks, Natick, MA, USA).

### 4.4 Assessment of the flow cell performance

To evaluate the rheometric performance of *bioRheoFlow*, measurements were performed on a benchmark PVA/borax solution and compared with results obtained using conventional geometries. Two stock solutions were prepared separately. PVA (*M*_*w*_ = 130 kDa) was dissolved at 4 wt% in 50 mL of Milli-Q water. The PVA powder was gradually added to the water to minimize agglomeration, and the solution was stirred overnight at 50 °C to 60 °C to promote complete dissolution. A separate 4 wt% borax solution was prepared by dissolving borax in 50 mL of Milli-Q water. The final PVA/borax solution was obtained by mixing 50 mL of the PVA stock solution with 10 mL of the borax stock solution, corresponding to a 5:1 v/v PVA-to-borax solution ratio. The mixture was stirred until a homogeneous viscoelastic material was obtained. Sample preparation followed established protocols for this material [54–56].

Oscillatory frequency sweeps obtained with *bioRheoFlow* were compared with measurements performed using conventional parallel-plate geometries with diameters of 10 mm and 25 mm and smooth surfaces, and a 20 mm roughened parallel-plate geometry with 100 µm-high pillars on both the upper and lower measuring surfaces intended to reduce wall slip.

Measurements were conducted at *T* = 25 °C, a gap of 1 mm, and a strain amplitude of *γ* = 1%, selected to remain within the linear viscoelastic regime. Dynamic frequency sweeps were performed from *ω* = 100 rads^*−*1^ to *ω* =0.1 rads^*−*1^ in the high-to-low frequency direction, using four measurement points per decade. A waiting time of 1 min was applied before each frequency sweep.

Instrument torque limits, inertia limits, and the onset of flow instabilities were calculated for each geometry and overlaid on the frequency-sweep data to indicate the operating windows, when relevant. Agreement between *bioRheoFlow* and the conventional geometries was assessed only within their common operating window. Excluding the high-frequency region affected by measuring-system inertia, the relative deviations in *G*^*′*^ and *G*^*′′*^ between the different geometries remained below 5%, indicating good agreement between *bioRheoFlow* and the conventional geometries.

Evaporation control was evaluated using oscillatory time sweeps at *ω* = 10 rads^*−*1^ and *γ* = 1%. Measurements were performed with *bioRheoFlow* and with conventional geometries operated with and without the manufacturer’s evaporation hood. For *bioRheoFlow*, Milli-Q water was continuously supplied through the lower porous plate at an infusion rate of 150 µLh^*−*1^ to compensate for solvent evaporation.

Each dynamic time sweep was performed for a fixed duration of 100 min and was not stopped before completion.

Solvent stability was quantified from the relative drift in the complex modulus |*G*^*\**^| with respect to its initial plateau value, with stable conditions defined as a variation in |*G*^*\**^| below 10% over the duration of the test. This comparison was used to assess the ability of *bioRheoFlow* to extend the stable measurement window relative to conventional geometries.

## Data availability

The research data supporting the findings of this study are available in the SigmaPlot file provided with the Supporting Information.

## Code availability

The scripts used for data analysis are available from the corresponding authors upon reasonable request.

## Acknowledgements

We acknowledge support of the project “Bioholistic: Developing integrated bioprocesses for a holistic chemical recycling of plastics” funded by the Swedish Research Council (FORMAS; Reference number 2022-00853) as the Lead Agency, Swiss National Science Foundation (SNSF) and Luxembourg National Research Fund (FNR) as the Partner Agencies, under the Weave cross-European initiative; ES acknowledges the support of the Swiss National Science Foundation (SNSF) (Grant No. 10005838).

## Author contributions

V.I.: Conceptualization, Methodology, Investigation, Formal analysis, Data curation, Software, Visualization, Writing – original draft.

S.G.: Methodology, Investigation.

E.S.: Conceptualization, Methodology, Supervision, Writing – review & editing.

T.T.: Supervision, Funding acquisition, Writing – review & editing.

J.V.: Conceptualization, Methodology, Supervision, Funding acquisition, Writing – review & editing.

## Supporting information

The Supporting Information includes technical drawings of the *bioRheoFlow* platform; details of tool assembly, alignment, parallelism, gap calibration, and measuring-system inertia; calibration of the liquid flow used to compensate for evaporation; representative observations and dynamic measurements of *in situ* biofilm growth under axial flow, including a bacteria-free control; the procedure used to identify the linear viscoelastic regime; the complete creep-test sequence for *Pantoea sp*.; the formulation and numerical implementation of the inertia-coupled Jeffreys model; and the evaluation of the rheometer operational windows, including minimum torque, modulus, and creep-compliance limits. The archive `bioRheoFlow_STL.zip` contains the STL files required to 3D print the *bioRheoFlow* components compatible with the rheometer used in this work.

## Supporting Information

### S1 Technical drawing of bioRheoFlow

Fig. S1 depicts the technical drawing of bioRheoFlow. In particular, Fig. S1a shows the bottom cover of the Peltier cell, while Fig. S1b shows the top and bottom casings (1 and 2, respectively) used as the measuring system on the rheometer.

#### Tool assembly and alignment

The porous filters were glued into the respective housings of the top and bottom casings, as described in the main text. Prior to alignment, the surfaces of the filters embedded in the resin-printed casings were flattened. To polish and level the top surface, a flexible J-weight cloth abrasive (sianor J, Sia Abrasives, Switzerland) with a P240 grit size was fixed onto the lower Peltier plate using double-sided adhesive tape. The top casing was brought into contact with the sandpaper under a normal force of approximately 1 N and rotated to achieve surface planarity. A similar procedure was applied to the bottom casing; in this case, the sandpaper was mounted on a standard 25 mm Anton Paar plate, and the same leveling procedure was performed (see Fig. S2).

Subsequently, the Peltier cover was installed and the leveling was verified using a spirit level placed at the center, as depicted in Fig. S3. The bottom casing was then fixed using three vertical screws.

The final step involved alignment of the top and bottom plates. This was performed visually with the aid of a camera to align the top and bottom plates.

In addition to leveling, the parallelism of the bottom geometry was assessed using a dial test indicator (TESATAST, TESA Technology, Hexagon), with a resolution of 0.002 mm.

As shown in Fig. S4, the dial indicator reading varies between 0.30 and 0.40 mm, indicating a deviation of approximately 0.10 mm.

Finally, the error in the gap (distance between top and bottom plates) after zero-gap calibration of the rheometer was evaluated using a feeler gauge (Spurtar, 32-blade set, 0.03 - 1 mm), resulting in a deviation of approximately 10 µm(see Fig. S5).

The last assessment of the new measuring system is the evaluation of the inertia of the tool. This final step can be performed by evaluating the “measuring system inertia” parameter in the rheometer software. The final values are reported in Table S1.

**Table S1:**
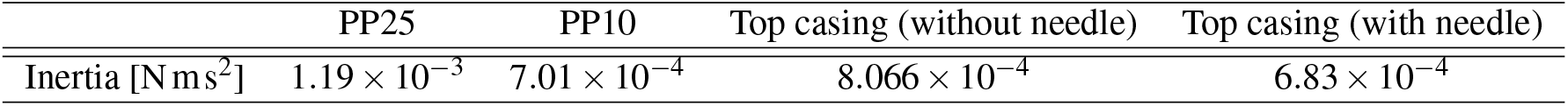
Measuring-system inertia values for the geometries used in the present work. The standard 10 mm and 25 mm plates are used as a reference.

|  | PP25 | PP10 | Top casing (without needle) | Top casing (with needle) |
| --- | --- | --- | --- | --- |
| Inertia [$\text{N m s}^2$] | $1.19 \times 10^{-3}$ | $7.01 \times 10^{-4}$ | $8.066 \times 10^{-4}$ | $6.83 \times 10^{-4}$ |

### S2 Calibrating the flow to counterbalance the evaporation

In the main manuscript, the tool presented in this work was initially employed to measure the rheology of biofilms grown externally and subsequently scraped into the rheometer measuring system. In order to counterbalance the drying of the biofilm during long rheological tests, we optimized the liquid flow supplied during the measurements. Figure S6 shows the evolution of the elastic modulus of a biofilm produced by *P. putida* when the flow is not properly balanced. This highlights the importance of precisely controlling the hydration conditions during biofilm rheology, since an incorrect balance can lead to significant artifacts in the measurements.

### S3 In situ growth under axial flow

Figure S7 shows a frame from the time-lapse video recording the biofilm growth during the first 24 hours, highlighting the main differences between the biofilms formed by *P. putida* and *Pantoea sp*. biofilms. *P. putida* forms a homogeneous biofilm front (Fig. S7a), whereas *Pantoea sp*. forms denser regions that float within the medium.

Figure S8 illustrates the increase in dynamic moduli during axial flow.

**Figure S1:**
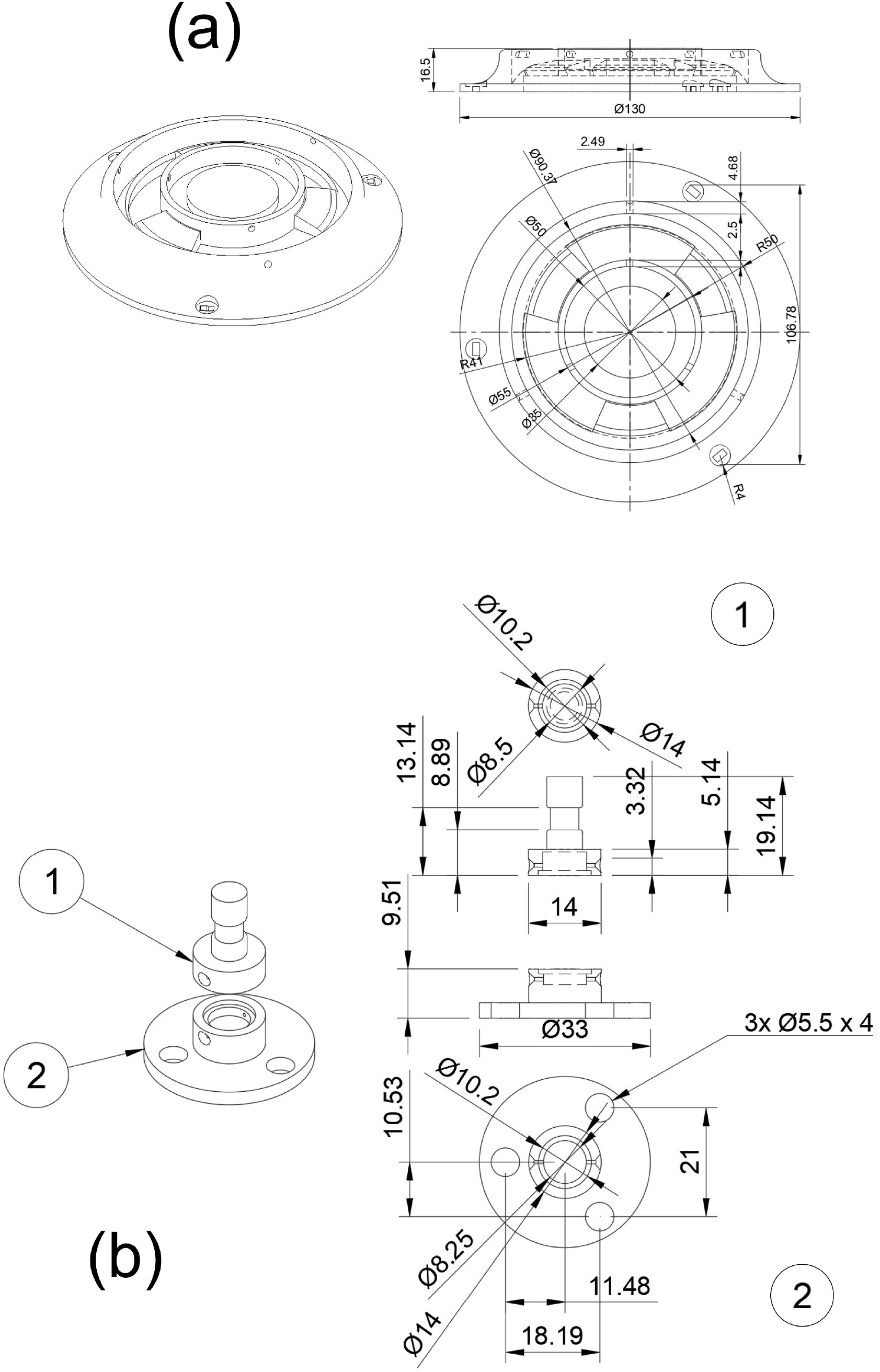
Technical drawing of the bioRheoFlow platform. (a) a base mount as a cover of the Peltier cell. (b) top (1) and bottom (2) casings

**Figure S2:**
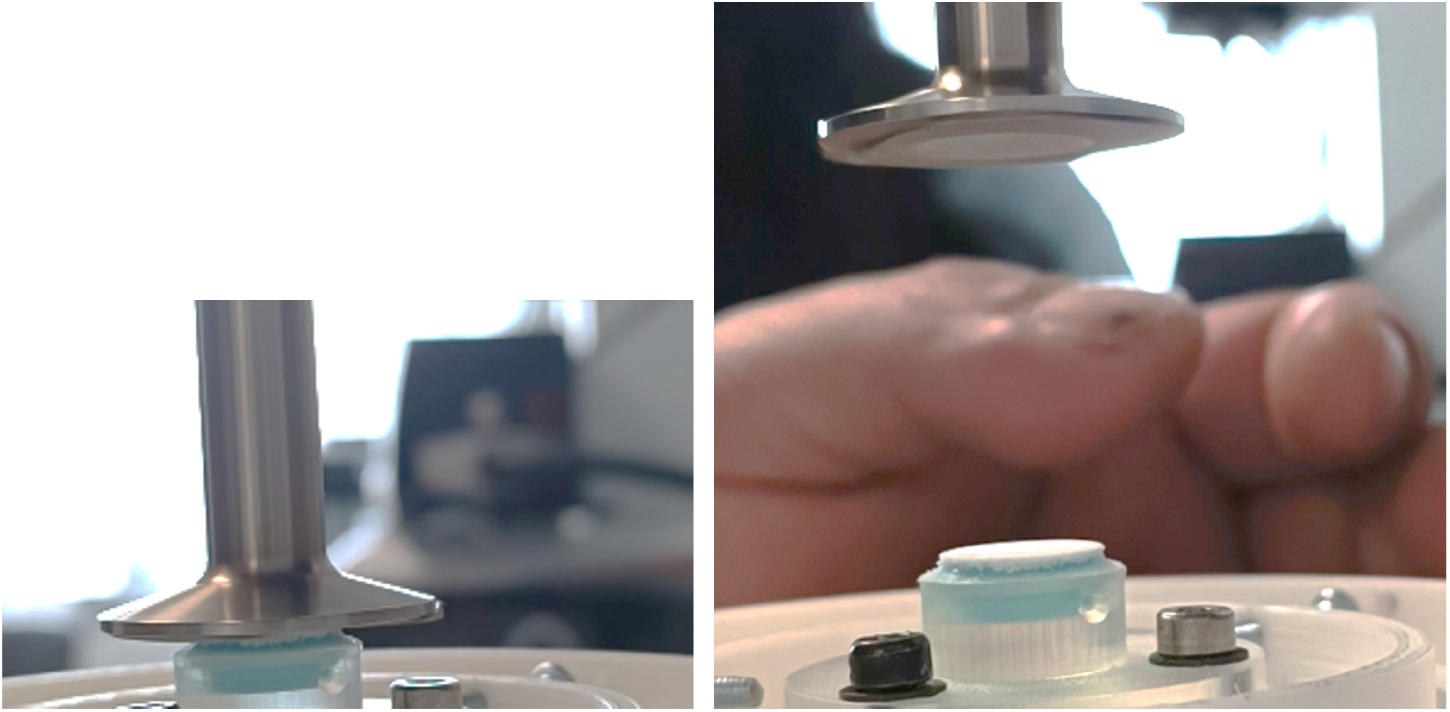
Standard PP25 geometry with attached double-sided adhesive tape.

**Figure S3:**
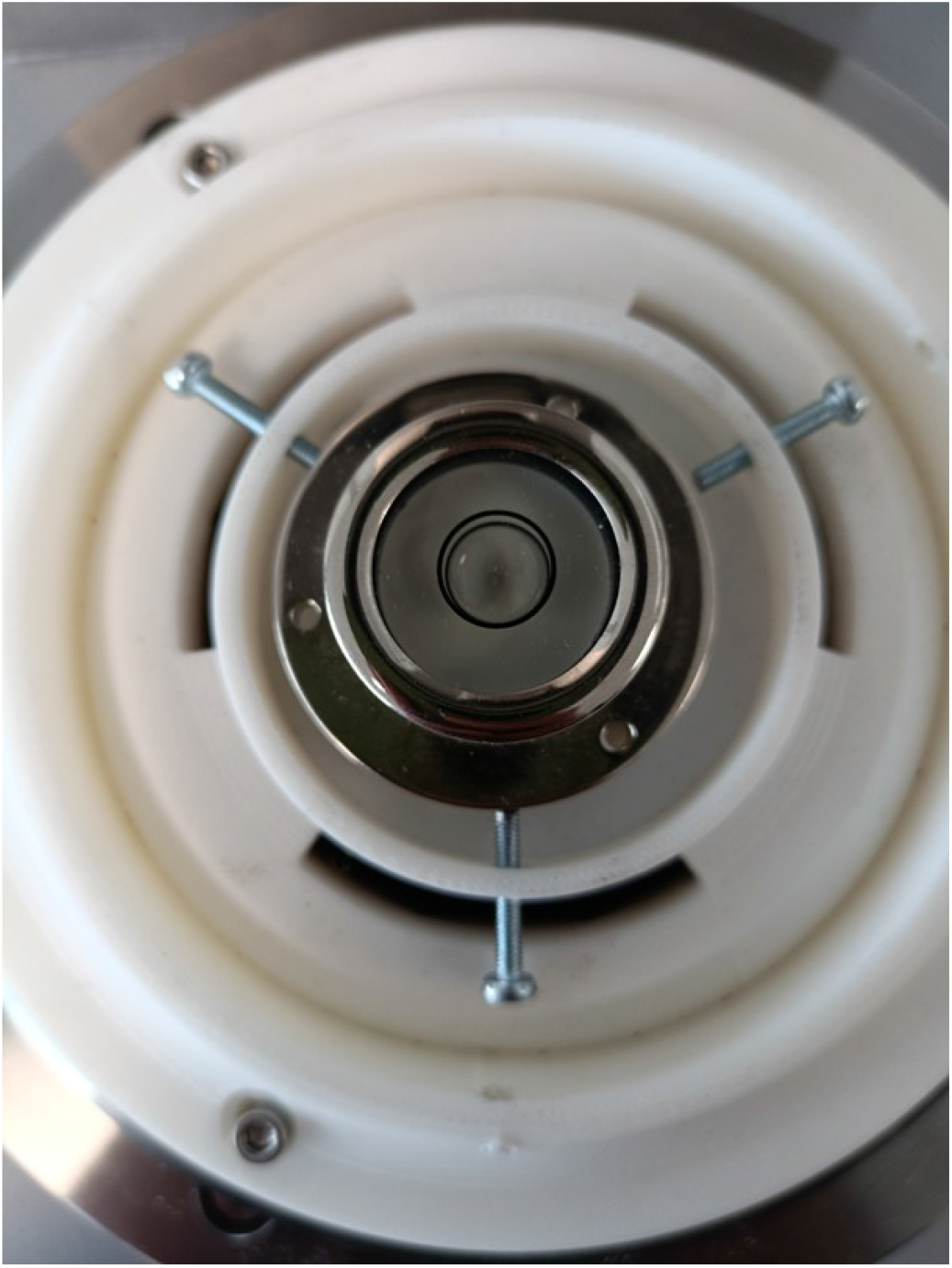
Cover of the Peltier with the spirit level in order to check the leveling of the tool.

**Figure S4:**
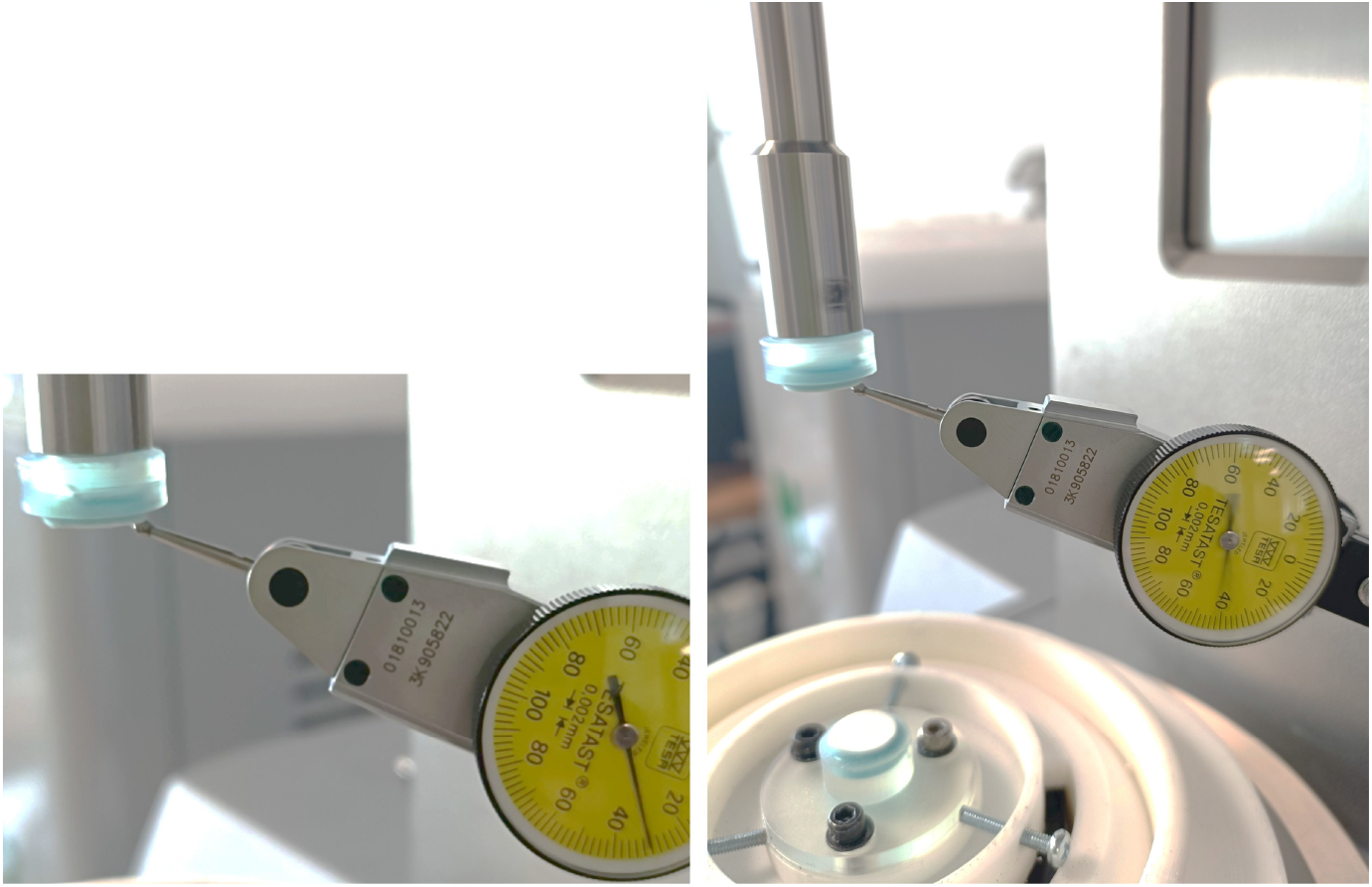
Dial test indicator. The tip is attached to the edge of the plate. The indicator oscillates between 0.3 and 0.4 mm

**Figure S5:**
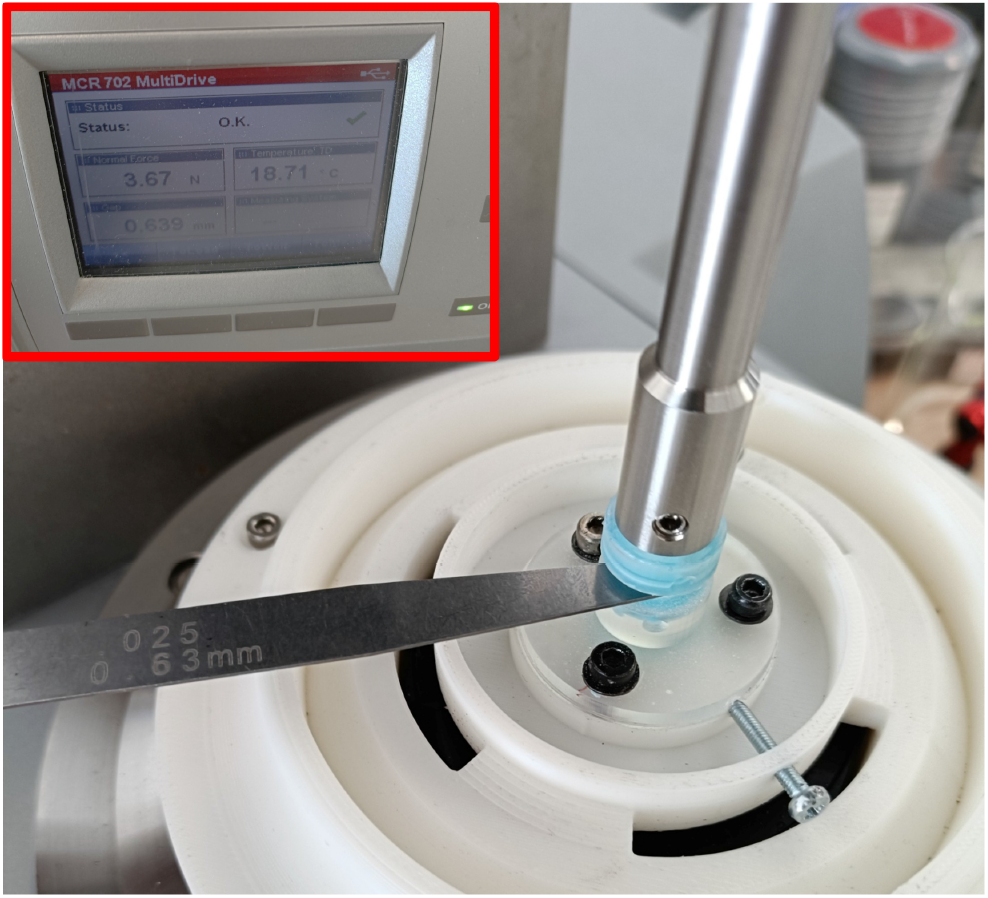
A feeler gauge with a thickness of 0.93 mm was used as reference. The rheometer registered a normal force of 3 N at a gap of 0.94 mm, corresponding to a distance error of approximately 0.01 mm.

**Figure S6:**
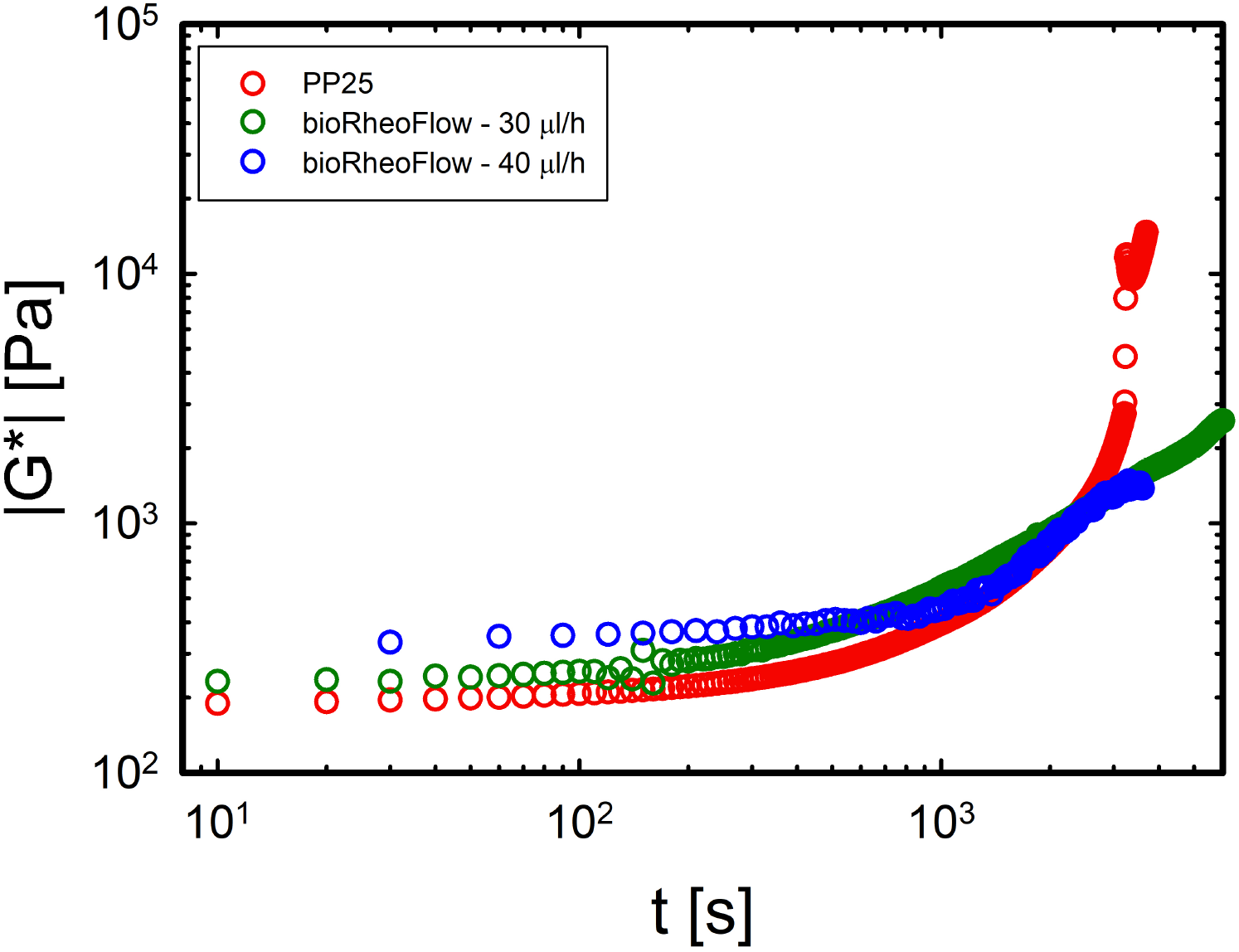
Comparison between a measurement performed with the conventional geometry (red circles) and two measurements performed with the proposed tool, where the medium flow does not counterbalance evaporation (blue and dark green circles).

Although further validation is needed, the results in Fig. S8 suggest that the growth, and thus the successful completion of the first stage, occurs when the modulus reaches a minimum, followed by a sharp increase. This criterion could therefore be used as an indicator for assessing the success of the initial growth phase.

The observed increase in the dynamic moduli is further supported by a control experiment conducted in the absence of bacteria. When the same protocol was applied to sterile LB medium, the dynamic moduli remained constant over time (see Fig. S9), supporting the interpretation that the increase observed in the presence of bacteria is associated with biofilm formation.

### S4 Evaluation of the linear viscoelastic regime

We report the procedure applied for the evaluation of the linear viscoelastic regime from the DSST. Data are reported as mean ± standard deviation from three independent biological replicates.

The dynamic moduli shown in Figures S10b and S10d are re-expressed in Figure S11 as functions of the shear stress.

Error bars are included on the x-axis to account for the variability and uncertainty inherent in stress values, which are calculated based on the applied strain. The dashed lines indicate the method used to determine the maximum shear stress value defining the limit of the linear viscoelastic regime (LVR). As in the previous cases, this threshold was established based on a 10% deviation in the elastic modulus. Analysis of the curves reveals that the LVR extends up to a critical shear stress of 9.61 ± 8.22 Pa for *P. putida* and 9.49 ± 6.49 Pa for *Pantoea sp*.. These critical stress values represent the mean and standard deviation calculated from the three independent biological replicates for each biofilm.

**Figure S7:**
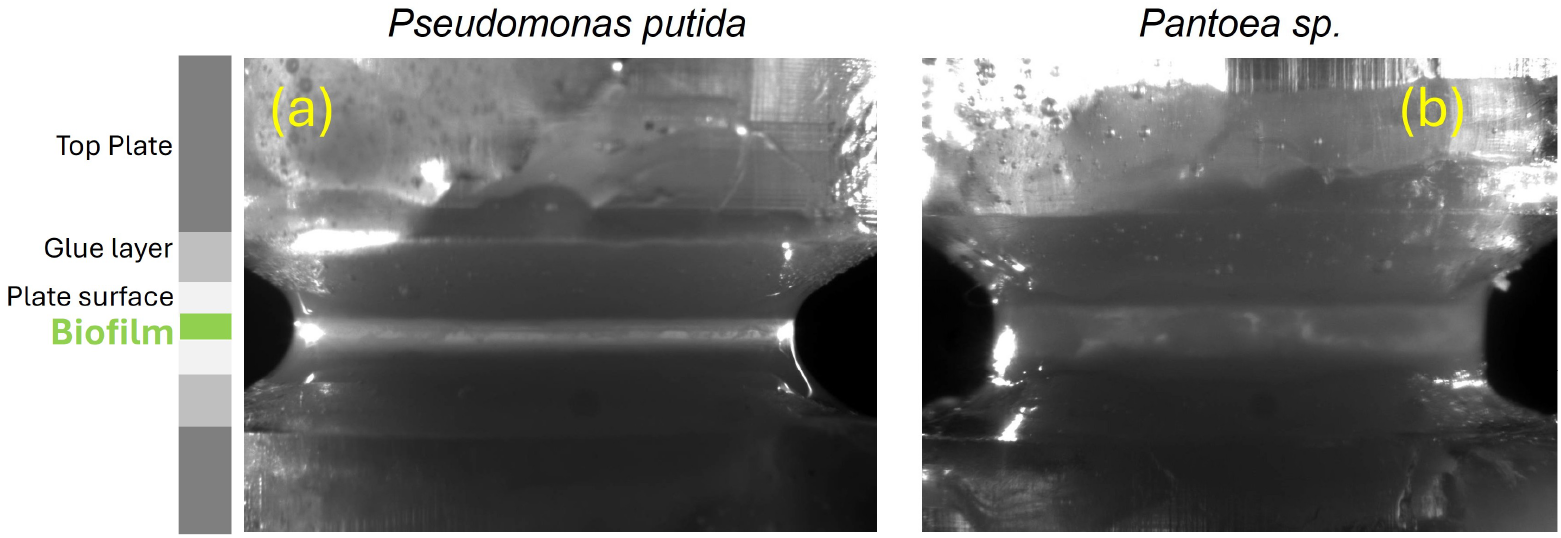
Representative frames acquired during *in situ* biofilm growth after ∼ 24 h under axial flow: (a) *P. putida*; (b) *Pantoea sp*.. The grayscale schematic identifies the components of the measuring geometry, with the biofilm position in the gap highlighted in green.

**Figure S8:**
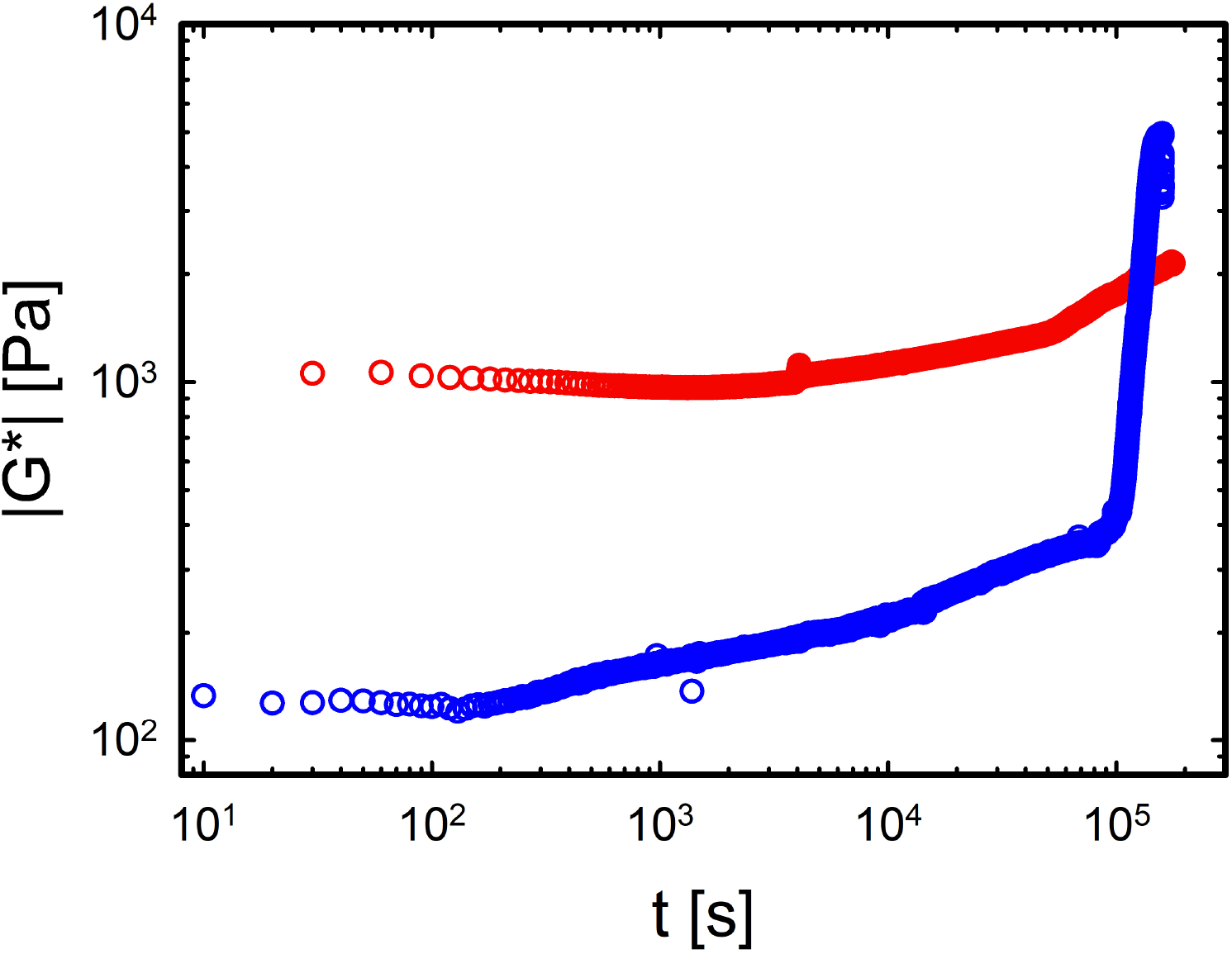
DTST during the in situ growth under the axial flow at *ω* =10 rads^*−*1^ for *P. putida* (red symbols) and *Pantoea sp*. (blue symbols). The measured moduli are reported as rheometric signals used to monitor the temporal evolution of the system during cultivation and are not interpreted as quantitative biofilm moduli because of the mechanical contribution of the axial-flow connection.

### S5 Creep tests on *Pantoea sp*

In Figure S12, the complete sequence of creep tests performed on *Pantoea sp*. is presented.

### S6 Jeffreys model

The analytical solution for the creep compliance, *J*(*t*), incorporating the rheometer’s rotational inertia *I* and the geometric form factor *b* (defined as *b* = *πR*^4^*/*2*h* for a parallel-plate geometry), can be expressed in terms of the Jeffreys model parameters. These material properties include the elastic modulus *G*_1_, the retardation viscosity *η*_1_ associated with the viscoelastic transient response, and the steady-state flow viscosity *η*_2_.

To simplify the mathematical formulation, the intermediate variables representing the damping factor, *A*, and the amplitude constant, *B*, are defined as follows:

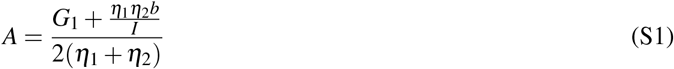

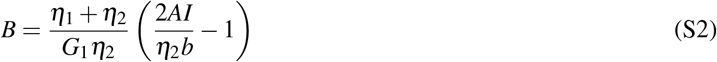

**Figure S9:**
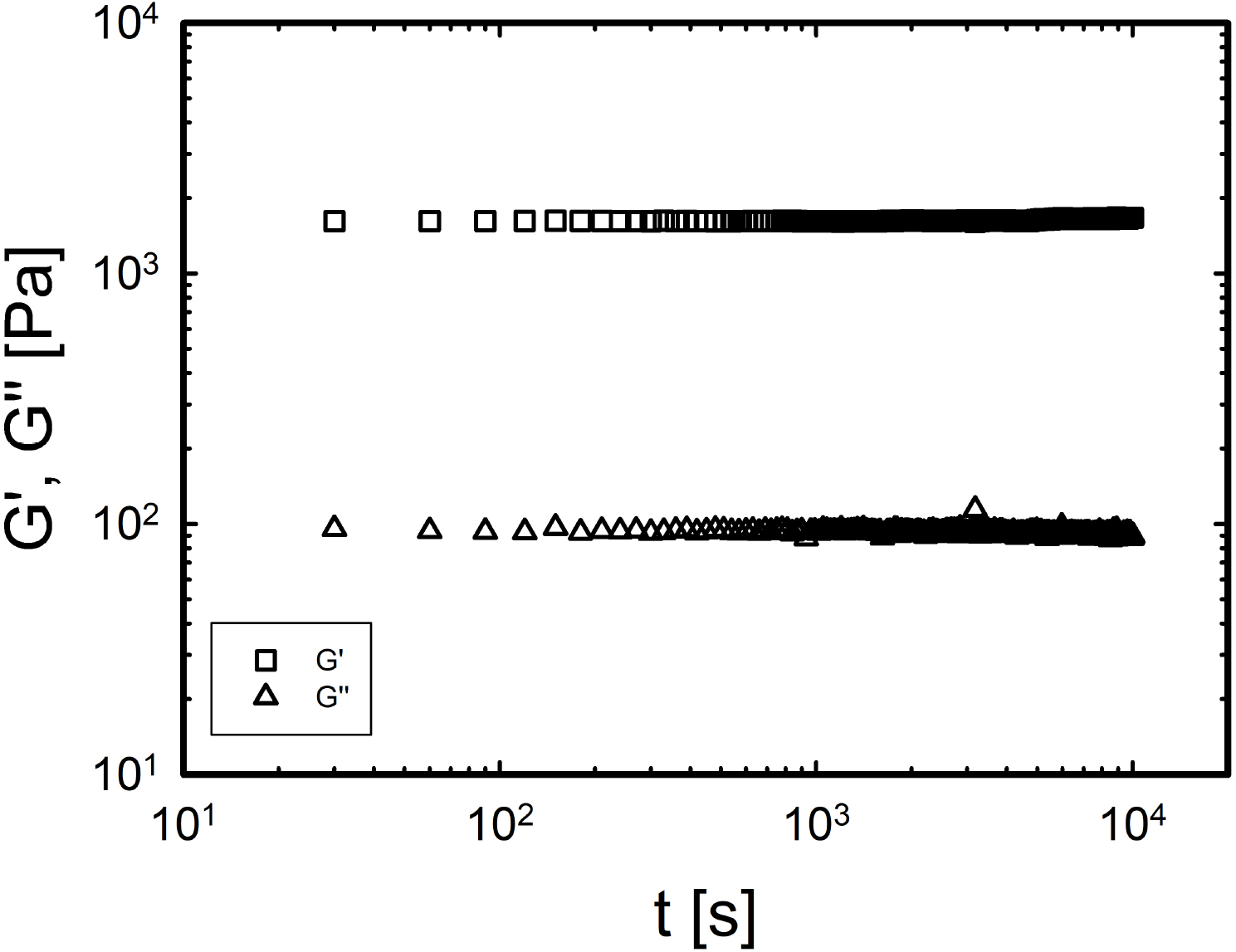
Dynamic time sweep test (DTST) during the axial flow of LB medium at *ω* = 10 rads^*−*1^ in the absence of bacteria. No increase in the moduli is observed, indicating the stability of the medium.

The angular frequency of the oscillatory response, *ω*, is derived as:

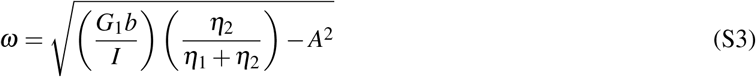

Finally, the full time-dependent creep compliance governing the coupled sample-instrument dynamics is expressed as:

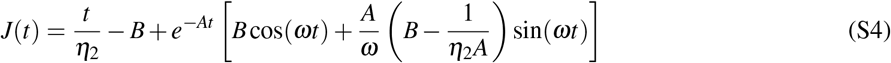

To extract the optimal material parameters (*G*_1_, *η*_1_, and *η*_2_), the theoretical model was fitted to the experimental creep compliance data by minimizing the objective function (*χ*^2^) defined in Eq.S5.

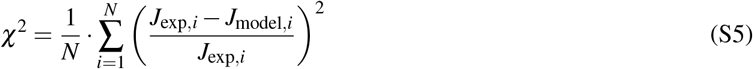

where *J*_exp_(*t*_*i*_) represents the experimentally measured compliance at time *t*_*i*_, *J*_model_(*t*_*i*_) is the theoretical compliance predicted by the model for the given set of parameters (*G*_1_, *η*_1_, *η*_2_), and *N* is the total number of experimental data points.

**Figure S10:**
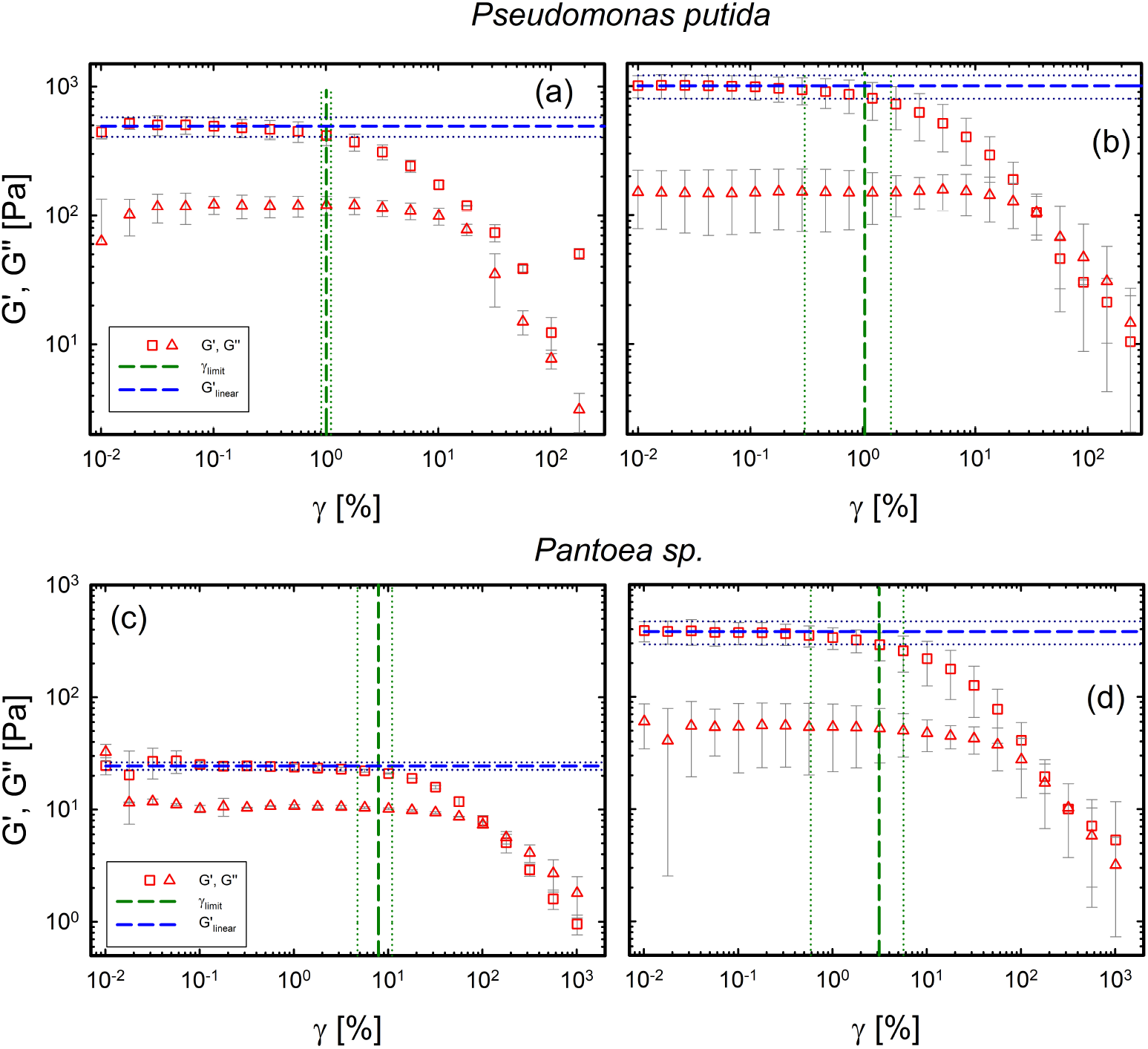
DSST (a), (c) on scraped biofilm of *P. putida* and *Pantoea sp*., respectively. DSST (b), (d) on biofilm grown under axial flow with our custom device of *P. putida* and *Pantoea sp*., respectively.

**Figure S11:**
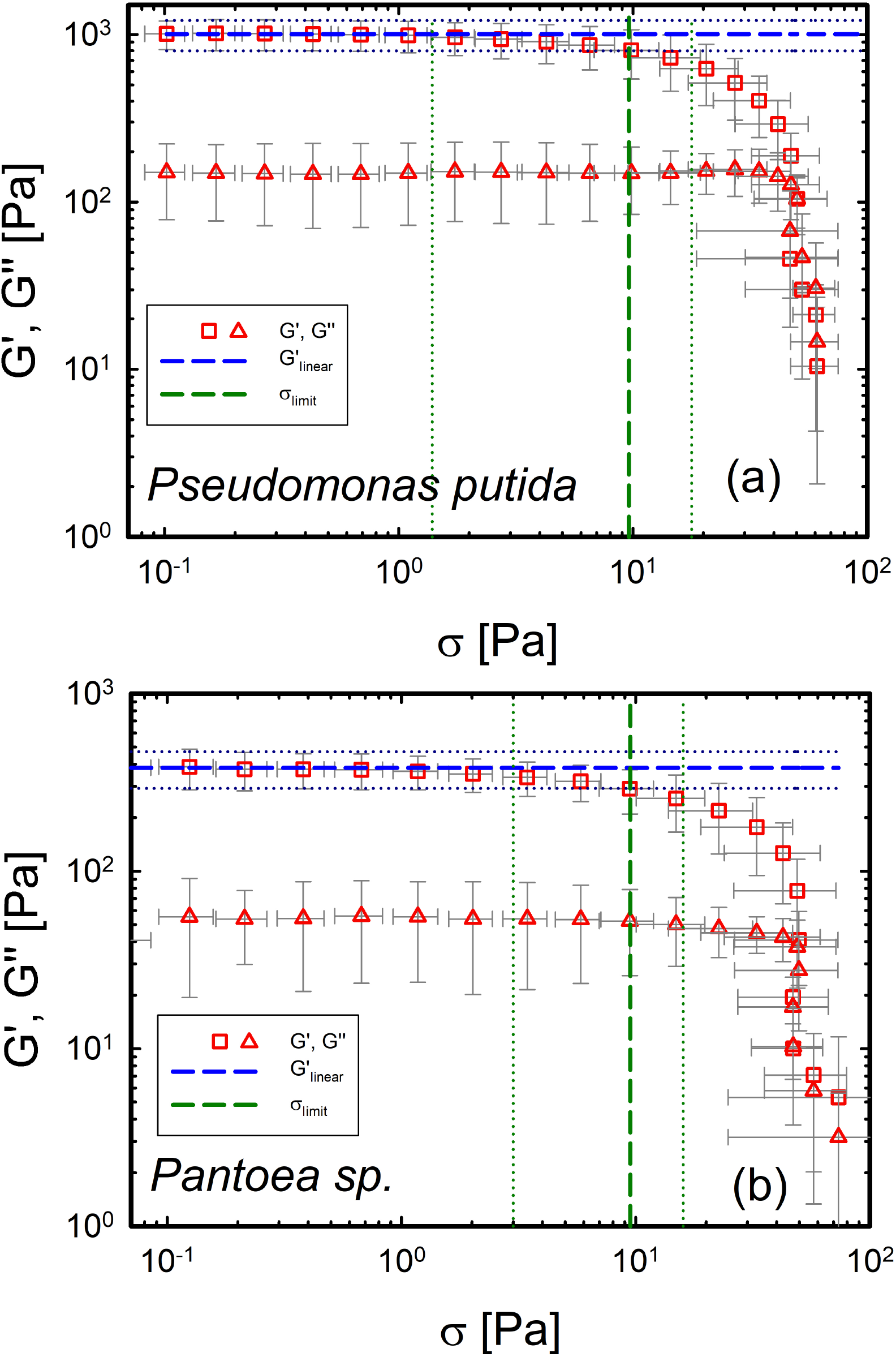
(a) DSST for *P. putida*. (b) DSST for *Pantoea sp*..

**Figure S12:**
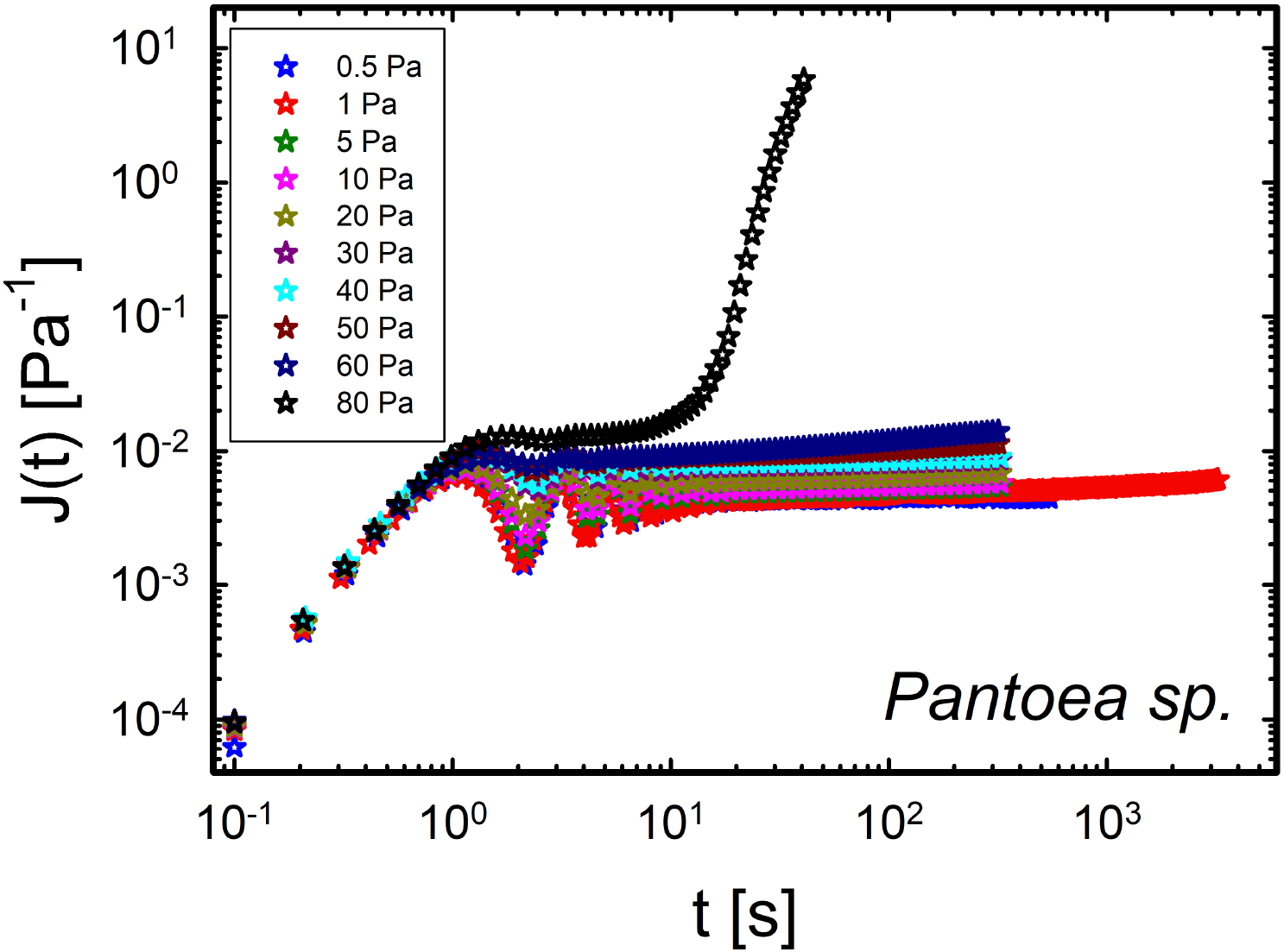
Creep tests on *Pantoea sp*.. The inset is a zoom on the behavior at an applied stress of 1 Pa, 5 Pa, 10 Pa.

To automatically extract the optimal material parameters, the minimization of the *χ*^2^ objective function was implemented using a custom MATLAB^®^ script. The numerical optimization was performed using the `fminsearchbnd` function based on the Nelder-Mead algorithm. Unlike `fminsearch`, which performs unconstrained optimization, `fminsearchbnd` allows the inclusion of lower and upper bounds. This was essential in our case to ensure that the parameters (*G*_1_, *η*_1_, and *η*_2_) remained strictly positive throughout the optimization, thus avoiding non-physical negative values of stiffness or viscosity.

### S7 Operational windows

In the main manuscript, the rheological results are presented together with the operational windows as defined by [43]. To adopt a more conservative approach to the reliability of the data, the evaluation of the minimum detectable modulus was based on the minimum torque sensitivity of the rheometer. According to the manufacturer (Anton Paar), the nominal minimum torque detectable in oscillatory mode is 0.5 nN·m and 1 nN·m in rotational. However, in this work, the minimum torque readable in oscillation was 6.95 nN·m and 19.7 nN·m in rotational. The calibration measurements, performed in both rotational and oscillatory modes to identify this minimum value, are shown in Fig. S13.

It is important to note that the minimum modulus reported herein does not refer to the complex modulus but specifically to the storage modulus, calculated as shown in Equation S7:

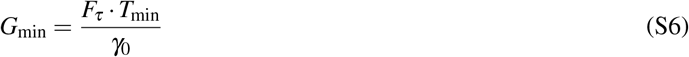

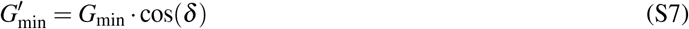

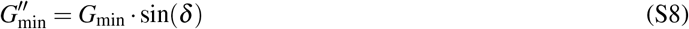

where *T*_min_ is the minimum torque applied, *γ*_0_ is the strain amplitude, *δ* is the phase angle in radians, and 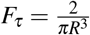 is the correction factor for torque for a plate-plate geometry.

**Figure S13:**
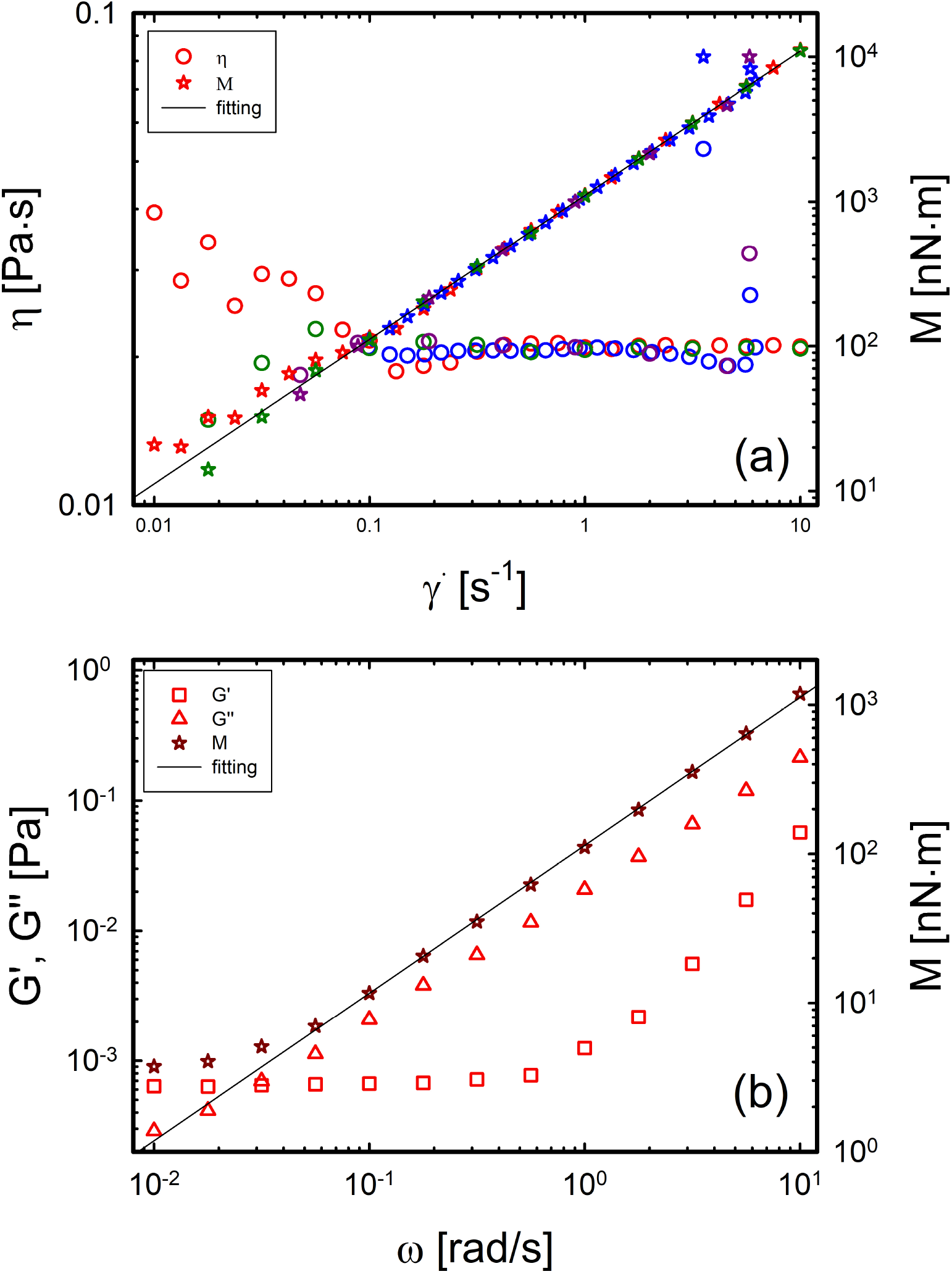
(a) Evaluation of the minimum torque during a steady measurements. (b) Evaluation of the minimum torque during a dynamic measurements.

Regarding the operational windows reported in Fig. 7, the minimum readable compliance can be derived from the definition of creep compliance:

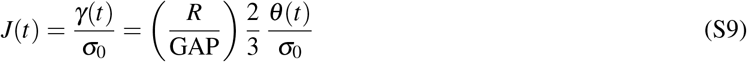

where *θ* is the deflection angle expressed in radians.

Considering the minimum applicable stress (*σ*_min_ = *F*_*τ*_ ·*M*_min_) and a minimum detectable angular displacement of *θ*_min_ = 10^*−*6^ rad, the minimum compliance can be expressed as:

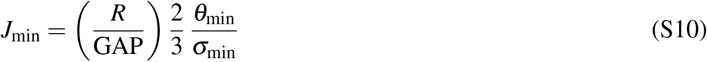

## References

(1) Nadell, C. D.; Xavier, J. B.; Foster, K. R. FEMS Microbiology Reviews 2009, 33, 206–224.

(2) Flemming, H. C.; Wuertz, S. Nature Reviews Microbiology 2019, 17, 247–260.

(3) Kochanowski, J. A.; Carroll, B.; Asp, M. E.; Kaputa, E. C.; Patteson, A. E. ACS Applied Bio Materials 2024, 7, 7809–7817.

(4) Flemming, H.-C.; Neu, T.; Wingender, J. The Perfect Slime: Microbial Extracellular Polymeric Substances (EPS) 2016, 1–14.

(5) Secchi, E. Physical Review E 2026, 114, 021001.

(6) Gloag, E. S.; Fabbri, S.; Wozniak, D. J.; Stoodley, P. Biofilm 2020, 2, 100017.

(7) Jeon, E.; Kim, H.; Kim, G.; Lee, D. Korea-Australia Rheology Journal 2023, 35, 267–278.

(8) Geisel, S.; Secchi, E.; Vermant, J. Interface Focus 2022, 12, DOI: 10.1098/rsfs.2022.0032.

(9) Tierra, G.; Pavissich, J. P.; Nerenberg, R.; Xu, Z.; Alber, M. S. Journal of The Royal Society Interface 2015, 12, 20150045.

(10) Boudarel, H.; Mathias, J.-D.; Blaysat, B.; Grédiac, M. npj Biofilms and Microbiomes 2018, 4, 17.

(11) Savorana, G.; Słomka, J.; Stocker, R.; Rusconi, R.; Secchi, E. Soft Matter 2022, 18, 3878–3890.

(12) Lieleg, O.; Caldara, M.; Baumgärtel, R.; Ribbeck, K. Soft Matter 2011, 7, 3307–3314.

(13) Kovach, K.; Davis-Fields, M.; Irie, Y.; Jain, K.; Doorwar, S.; Vuong, K.; Dhamani, N.; Mohanty, K.; Touhami, A.; Gordon, V. D. Biofilms and Microbiomes 2017, 3, DOI: 10.1038/s41522-016-0007-9.

(14) Gloag, E. S.; German, G. K.; Stoodley, P.; Wozniak, D. J. Scientific Reports 2018, 8, 1–11.

(15) Kretschmer, M.; Schüßler, C. A.; Lieleg, O. Advanced Materials Interfaces 2021, 8, DOI: 10.1002/admi.202001658.

(16) Yan, J.; Moreau, A.; Khodaparast, S.; Perazzo, A.; Feng, J.; Fei, C.; Mao, S.; Mukherjee, S.; Košmrlj, A.; Wingreen, N. S.; Bassler, B. L.; Stone, H. A. Advanced Materials 2018, 30, DOI: 10.1002/adma.201804153.

(17) Klotz, M.; Kretschmer, M.; Goetz, A.; Ezendam, S.; Lieleg, O.; Opitz, M. RSC Advances 2019, 9, 11521–11529.

(18) Horvat, M.; Pannuri, A.; Romeo, T.; Dogsa, I.; Stopar, D. Soft Matter 2019, 15, 5042–5051.

(19) Houari, A.; Picard, J.; Habarou, H.; Galas, L.; Vaudry, H.; Heim, V.; Di Martino, P. Biofouling 2008, 24, 235–240.

(20) Kesel, S.; Grumbein, S.; Gümperlein, I.; Tallawi, M.; Marel, A. K.; Lieleg, O.; Opitz, M. Applied and Environmental Microbiology 2016, 82, 2424–2432.

(21) Romero, C. M.; Martorell, P. V.; López, A. G.; Peñalver, C. G.; Chaves, S.; Mechetti, M. Colloids and Surfaces B: Biointerfaces 2018, 162, 246–255.

(22) Charlton, S. G.; White, M. A.; Jana, S.; Eland, L. E.; Jayathilake, P. G.; Burgess, J. G.; Chen, J.; Wipat, A.; Curtis, T. P. Journal of Bacteriology 2019, 201, DOI: 10.1128/JB.00101-19.

(23) Jana, S.; Charlton, S. G.; Eland, L. E.; Burgess, J. G.; Wipat, A.; Curtis, T. P.; Chen, J. Biofilms and Microbiomes 2020, 6, DOI: 10.1038/s41522-020-0126-1.

(24) Siri, M.; Sarlet, A.; Ziege, R.; Zorzetto, L.; Sotelo Guzman, C.; Amini, S.; Hengge, R.; Blank, K. G.; Bidan, C. M. ACS Biomaterials Science & Engineering 2025, 11, 4523–4536.

(25) Sato, J.; Breedveld, V. Applied Rheology 2005, 15, 390–397.

(26) Magnin, A.; Piau, J. Journal of Non-Newtonian Fluid Mechanics 1990, 36, 85–108.

(27) Orafidiya, L. O. Journal of pharmacy and pharmacology 1989, 41, 341–342.

(28) Nommensen, P.; Duits, M.; Van Den Ende, D.; Mellema, J. Physical Review E 1999, 59, 3147.

(29) Louhichi, A.; Jacob, A.; Bouteiller, L.; Vlassopoulos, D. Journal of Rheology 2017, 61, 1173–1182.

(30) Giacomin, A. J.; Pasquino, R.; Saengow, C.; Gilbert, P. H. Rheologica Acta 2019, 58, 291–305.

(31) Pavlovsky, L.; Younger, J. G.; Solomon, M. J. Soft matter 2013, 9, 122–131.

(32) Waters, M. S.; Kundu, S.; Lin, N. J.; Lin-Gibson, S. ACS applied materials & interfaces 2014, 6, 327–332.

(33) Yannarell, S. M.; Grandchamp, G. M.; Chen, S.-Y.; Daniels, K. E.; Shank, E. A. Journal of bacteriology 2019, 201, 10–1128.

(34) Rusconi, R.; Lecuyer, S.; Autrusson, N.; Guglielmini, L.; Stone, H. A. Biophysical Journal 2011, 100, 1392–1399.

(35) Rusconi, R.; Lecuyer, S.; Guglielmini, L.; Stone, H. A. Journal of the Royal Society Interface 2010, 7, 1293–1299.

(36) Purevdorj, B.; Costerton, J. W.; Stoodley, P. Applied and Environmental Microbiology 2002, 68, 4457–4464.

(37) Drescher, K.; Shen, Y.; Bassler, B. L.; Stone, H. A. Proceedings of the National Academy of Sciences 2013, 110, 4345–4350.

(38) Krsmanovic, M.; Biswas, D.; Ali, H.; Kumar, A.; Ghosh, R.; Dickerson, A. K. Advances in Colloid and Interface Science 2021, 288, DOI: 10.1016/j.cis.2020.102336.

(39) Conrad, J. C.; Poling-Skutvik, R. Annual Review of Chemical and Biomolecular Engineering 2018, 9, 175–200.

(40) Tsagkari, E.; Connelly, S.; Liu, Z.; McBride, A.; Sloan, W. T. npj Biofilms and Microbiomes 2022, 8, DOI:10.1038/s41522-022-00300-4.

(41) Stoodley, P.; Cargo, R.; Rupp, C. J.; Wilson, S.; Klapper, I. Journal of Industrial Microbiology and Biotechnology 2002, 29, 361–367.

(42) Galy, O.; Latour-Lambert, P.; Zrelli, K.; Ghigo, J. M.; Beloin, C.; Henry, N. Biophysical Journal 2012, 103, 1400–1408.

(43) Ewoldt, R. H.; Johnston, M. T.; Caretta, L. M. In Complex Fluids in Biological Systems, Spagnolie, S. E., Ed.; Springer: 2014, pp 207–241.

(44) Mewis, J.; Wagner, N. J., Colloidal suspension rheology; Cambridge university press: 2012.

(45) Ewoldt, R.; McKinley, G. H. Rheol. Bull 2007, 76.

(46) Towler, B. W.; Rupp, C. J.; Cunningham, A. B.; Stoodley, P. Biofouling 2003, 19, 279–285.

(47) Goudoulas, T. B.; Germann, N. Journal of Rheology 2016, 60, 491–502.

(48) Baravian, C.; Quemada, D. Rheologica acta 1998, 37, 223–233.

(49) Flory, P. J., Principles of polymer chemistry; Cornell university press: 1953.

(50) Dodero, A.; Williams, R.; Gagliardi, S.; Vicini, S.; Alloisio, M.; Castellano, M. Carbohydrate Polymers 2019, 203, 349–355.

(51) Borzacchiello, A.; Russo, L.; Malle, B. M.; Schwach-Abdellaoui, K.; Ambrosio, L. BioMed research international 2015, 2015, 871218.

(52) Puhm, M.; Ainelo, H.; Kivisaar, M.; Teras, R. Microorganisms 2022, 10, 618.

(53) Charlton, S. G.; Bible, A. N.; Secchi, E.; Morrell-Falvey, J. L.; Retterer, S. T.; Curtis, T. P.; Chen, J.; Jana, S. Advanced Science 2023, 10, 2207373.

(54) McLaughlin, K. W.; Wyffels, N. K.; Jentz, A. B.; Keenan, M. V. Journal of Chemical Education 1997, 74, 97.

(55) Mahjoub, H. F.; Zammali, M.; Abbes, C.; Othman, T. Journal of Molecular Liquids 2019, 291, 111272.

(56) Casassa, E.; Sarquis, A.; Van Dyke, C. Journal of Chemical Education 1986, 63, 57.

